# Untangling Microbiota Diversity and Assembly Patterns in the World’s Largest Water Diversion Canal

**DOI:** 10.1101/2021.07.26.453773

**Authors:** Lu Zhang, Wei Yin, Chao Wang, Aijing Zhang, Hong Zhang, Tong Zhang, Feng Ju

## Abstract

Large water diversion projects are important constructions for reallocation of human-essential water resources. Deciphering microbiota dynamics and assembly mechanisms underlying canal water ecosystem services especially during long-distance diversion is the prerequisite for water quality monitoring, biohazard warning and sustainable management. Using a 1432-km canal of the South-to-North Water Diversion Projects as a model system, we answer three central questions: how bacterial and micro-eukaryotic communities spatio-temporally develop, how much ecological stochasticity contributes to microbiota assembly, and which immigrating populations better survive and navigate across the canal. We applied quantitative ribosomal RNA gene sequence analyses to investigate canal water microbial communities sampled over a year, as well as null model- and neutral model-based approaches to disentangle the microbiota assembly processes. Our results showed clear microbiota dynamics in community composition driven by seasonality more than geographic location, and seasonally dependent influence of environmental parameters. Overall, bacterial community was largely shaped by deterministic processes, whereas stochasticity dominated micro-eukaryotic community assembly. We defined a local growth factor (LGF) and demonstrated its innovative use to quantitatively infer microbial proliferation, unraveling taxonomically dependent population response to local environmental selection across canal sections. Using LGF as a quantitative indicator of immigrating capacities, we also found that most micro-eukaryotic populations (82%) from the source lake water sustained growth in the canal and better acclimated to the hydrodynamical water environment than bacteria (67%). Taxa inferred to largely propagate include *Limnohabitans* sp. and *Cryptophyceae*, potentially contributing to water auto-purification. Combined, our work poses first and unique insights into the microbiota assembly patterns and dynamics in the world’s largest water diversion canal, providing important ecological knowledge for long-term sustainable water quality maintenance in such a giant engineered system.

## Introduction

Microorganisms are fundamental components of ecosystems and play vital roles in global biogeochemical cycling (Arrigo, 2005; Battin et al., 2016; Du et al., 2021). Knowledge about composition and dynamics of microbial communities across space and time is a prerequisite for predicting and manipulating the microorganisms in both natural and engineered systems (Bell et al., 2005; Bru et al., 2011; Ju and Zhang, 2015). Due to artificial system design and control, microbial communities in engineered ecosystems have received increasing attention to unravel the community dynamics underlying their eco-environmental or biotechnological services (Briones and Raskin, 2003; Ju et al., 2017; Ju et al., 2014; Vanwonterghem et al., 2014; Wu et al., 2019). Specifically, microorganisms in drinking water systems are closely linked with water auto-purification and eventually to public health (Berry et al., 2006). Thus, comprehensive microbiome monitoring is needed to differentiate baseline and disturbed microbial communities in engineered water systems to facilitate system management (Hull et al., 2019; Sagova-Mareckova et al., 2021).

The South-to-North Water Diversion (SNWD) Projects were built to aid in water supply in the drought-stricken north China, which represent the world’s largest water diversion projects. The projects consist of the Eastern Route Project, the Middle Route Project (MRP) and the Western Route Project. The 1432-km MRP went into service in 2014 and has drawn 40 billion m^3^ of water from the Danjiangkou Reservoir to He’nan and Hebei Provinces, and Beijing and Tianjin Municipalities, benefiting tens of millions of people. Despite their important roles in controlling geochemical processes and indicating water quality (Chen et al., 2018; Dang et al., 2018), little has been known about the microbial constituents throughout the system (Luo et al., 2019), neither for other engineered long-distance water diversion canals. Especially, the MRP is a closed engineered system where microorganisms are subject to potentially less complex biotic and abiotic interactions, while the numerous hydraulic structures and engineered perturbations (e.g., inverted siphon, aqueduct and tunnels) along the main canal add to challenges to model microbial community dynamics, calling for comprehensive investigation of microbial community patterns in the MRP canal, to assist in establishing integrative assessment of water quality with microbial community monitoring (Sagova-Mareckova et al., 2021).

Understanding processes and mechanisms driving the community patterns underlying ecosystem services is one of the key issues in microbial ecology (Zhou et al., 2014). Microbial community composition in aquatic ecosystems can be affected deterministically by both abiotic or environmental factors, e.g., water temperature, pH, dissolved oxygen, and nutrient availability (Hu et al., 2017; Liu et al., 2020a; Zhang et al., 2019b), and biotic factors, such as competition and predation. In addition to deterministic mechanisms, stochastic processes, for instance, dispersal and ecological drift, also contribute to microbial community assembly. Efforts have been made to disentangle the relative importance of deterministic vs. stochastic processes based on a variety of null and neutral modeling methods (Ning et al., 2019; Sloan et al., 2006; Stegen et al., 2013). Depending on which ecosystems or organism types these methods are applied to, the relative importance of deterministic vs. stochastic processes may greatly vary (Chen et al., 2019; Liu et al., 2018b; Wang et al., 2020; Yan et al., 2020; Niederdorfer et al., 2021). For instance, bacterial communities in lake water were revealed to be predominantly structured by determinism, whereas micro-eukaryotic communities largely by stochasticity (Logares et al., 2018). For the MRP, the closed channel, the environmental gradients and the variety of hydraulic constructions along the canal make it obscure to what extent ecological stochasticity determines the assembly of microbiota community, which is essential for an improved understanding of the ecosystem processes and microbial dynamics prediction.

Other emerging questions in such large water-diversion canal and the comparable systems, like river continuum (Gweon et al., 2021), include which immigrating microorganisms are adapted to local niches in the receiving water and how they functionally contribute to the ecological service. Especially in the MRP, how many of the microbial populations originated from a natural reservoir may survive in the contrastingly engineered canal remains an open question. Although population growth associated with microbial immigration has been quantified in other engineered water systems (Mei and Liu, 2019), such as activated sludge reactors (Mei et al., 2019), efforts in estimating microbial growth in long-distance and evolving systems are still lacking.

In the present study, we followed the spatio-temporal dynamics of prokaryotic and micro-eukaryotic community along the main canal of MRP of the SNWD project over a year to address: 1) How does microbial community develop with the water flow? 2) How is the relative importance of deterministic and stochastic processes on shaping the community assembly in this closed turbulent engineered system? 3) How do the immigrating bacteria and micro-eukaryotes from water source survive, grow, or die off in the canal? Based on high-throughput marker gene (i.e., 16S and 18S) amplicon and metagenomic sequencing, bioinformatics, and statistical modeling of water microbiota, we elucidated the microbial diversity and population dynamics along the canal over four seasons and uncovered the key patterns and processes driving community assembly. By overcoming the limitations of widely used relative abundance measures with estimated absolute abundance (Barlow et al., 2020; Tkacz et al., 2018), we demonstrated the first use of local growth factor to predict growth capacity of the immigrating microbial populations along the water canal and revealed the divergent microbial fates and population lifestyles throughout the world’s largest water diversion canal.

## Materials and Methods

A full version of the Materials and methods are available in the Supplementary Information.

### Study sites, sampling and physiochemical monitoring

The Middle Route Project (MRP) of the South-to-North Water Diversion (SNWD) Projects started from Danjiangkou reservoir and diverted water northwards until Beijing and Tianjin, which is an artificial system without direct connection with surface waters in the area. Fifty samples were collected from 19 sites (Fig. 1), including one at the Danjiangkou reservoir (S01) and 18 along the main canal of the MRP (S02-S18). The 18 sampling sites along the canal were selected to capture the spatial dynamics of microbial communities from canal head to end across the subtropical and warm-temperate climate zones. The sites were grouped to three sections: S02-S08 (Section1), S09p-S14 (Section2) and S15-S18 (Section3). Canal water was sampled and filtered through 0.22-μm cellulose nitrate filters (Sartorius, Germany). Physiochemical properties, including water temperature (T), pH, dissolved oxygen (DO), total nitrogen (TN) and fluoride (F^-^), were measured according to the environmental quality standard for surface water of China (GB3838-2002). Details about the sampling sites and physiochemical measurements are in the Supplementary Method S1.

**Fig. 1.**
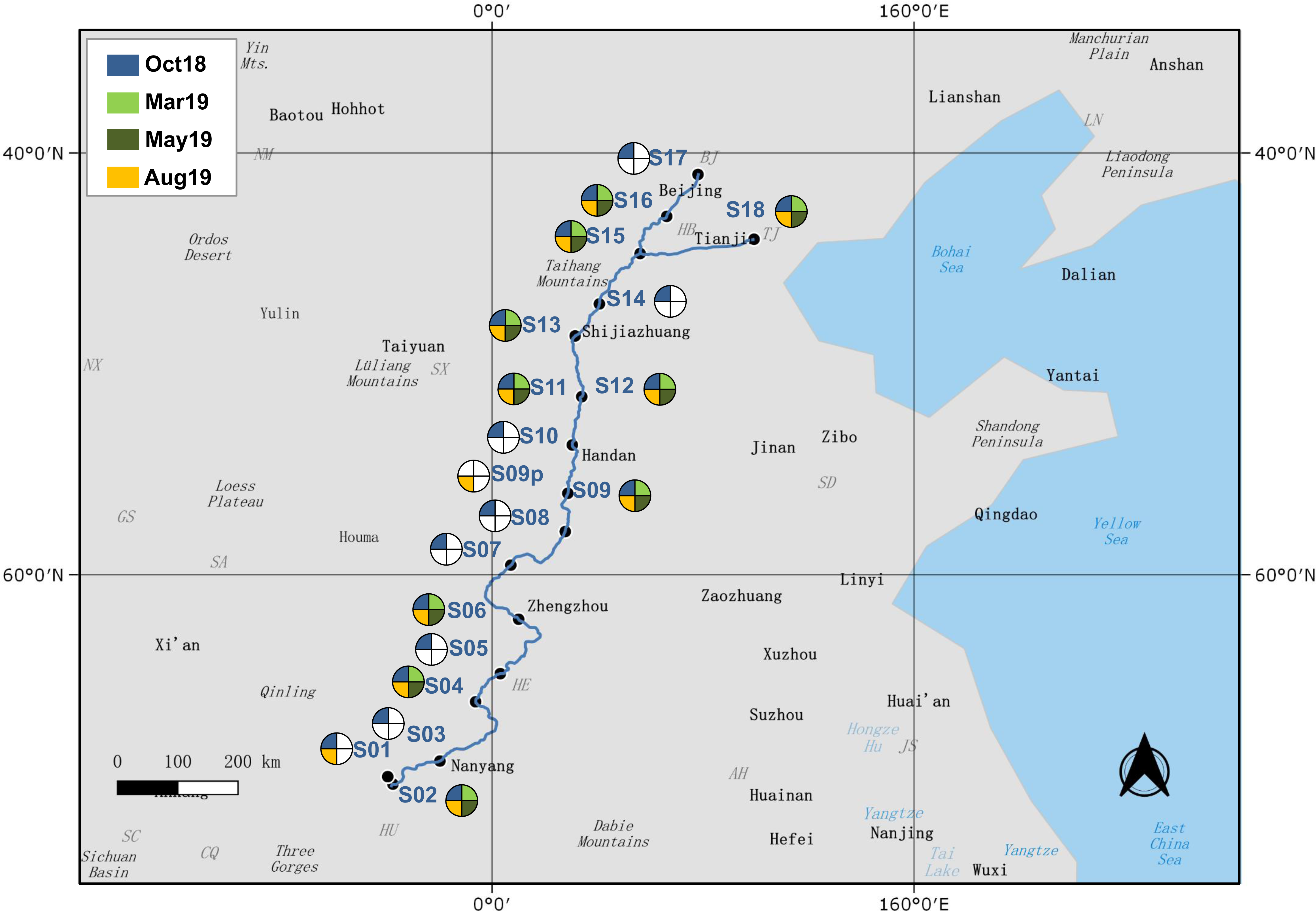
Four sampling campaigns at 19 sites along the 1432-km main canal of the MRP of the SNWD project selected for microbial and physicochemical analyses. Colored fans indicate months of sampling at each site. The map was created using QGIS (version 3.10.5, www.qgis.org).

### DNA extraction, PCR amplification and sequencing

Genomic DNA was extracted from filters using FastDNA Spin Kit for Soil (MP Biomedicals, USA). Bacterial 16S rRNA gene V3-V4 region and eukaryotic 18S rRNA gene V4 region were amplified and sequenced on the Illumina platforms producing 250 bp paired-end reads at the Guangdong Magigene Biotechnology Co., Ltd. (Guangzhou, China). Real-time quantitative PCR (qPCR) was additionally performed to quantify the 16S rRNA gene copies (as a proxy for bacterial biomass) in each genomic DNA sample, as described in Ju et al. (2019). Detailed methodologies are provided in the Supplementary Method S2.

Each genomic DNA was also used for shot-gun metagenomic sequencing on the Illumina’s Novaseq platform using a 150 bp paired-end sequencing strategy at the Novogene (Beijing, China). The sequencing data generated in this study have been deposited into CNGB Sequence Archive (CNSA) (Guo et al., 2020) of China National GeneBank DataBase (CNGBdb) (Chen et al., 2020) with accession number CNP0001593.

### Bioinformatics and statistical analyses of microbial diversity

The 16S and 18S rRNA gene amplicon sequence data were processed using Qiime2 pipeline (Bolyen et al., 2019) and DADA2 algorithm (Callahan et al., 2016) as described in the Supplementary Method S3, generating final abundance tables of amplicon sequence variants (ASVs) consisting of 6705 bacterial and 3217 eukaryotic ASVs, respectively. The alpha-diversity metrics of bacterial and micro-eukaryotic communities were further computed in Qiime2. All the univariate and multivariate statistics were performed in R (R Core Team, 2020). A set of spatial factors was generated for the sampling sites based on the geographic coordinates using the principal coordinates of neighboring matrices (PCNM) analysis (Borcard and Legendre, 2002).

The metagenomic data were pretreated and searched against the SILVA SSU132 database to identify rRNA gene reads. This information was further used to estimate the 16S rRNA and 18S rRNA gene copies detected in each metagenome based on the sum for the read coverage over the full length of each matched reference sequence in the database. Equation and details are available in the Supplementary Method S3. The copies of 18S rRNA genes in each water sample were estimated as 16S rRNA gene copies (determined by the aforementioned qPCR analysis) divided by ratios of 16S rRNA to 18S rRNA gene copy (Table S1) as sequenced and estimated in the metagenomic data.

### Neutral and null model analyses of microbial community patterns

To disentangle the relative contribution of stochastic and deterministic processes to microbial community assembly, Sloan neutral model (Sloan et al., 2006) and null model-based NST approach (Ning et al., 2019) were applied to the 16S and 18S community data. Fitting of the model were performed in R according to Burns et al. (2016). Accordingly, ASVs were classified into three groups depending on whether their observed occurrence falling higher than (“above prediction”), within (“as predicted”) or lower than (“below prediction”) the 95% confidence interval of the neutral model predictions. Normalized stochasticity ratio (NST) was used to quantify ecological stochasticity in communities within canal sections (Ning et al., 2019). NST analysis was performed in R using the NST package (Ning et al., 2019). More details about the methodology are in the Supplementary Method S4.

### Population lifestyle and local growth factor analyses

The absolute abundance of bacterial and micro-eukaryotic ASVs was calculated by multiplying relative abundance of each ASV computed from amplicon sequence data by the overall 16S and 18S rRNA gene copies detected in a given metagenome for all the Oct samples (see the Supplementary Method S5 for more details).

To characterize fate of microorganisms originated from natural reservoir after transported into the closed canal, we defined a local growth factor (LGF) to depict whether the populations grew or restrained, by comparing their mean absolute abundance in each canal section to the abundance at the canal head, as follows:

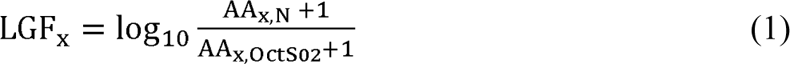

where AA_x, N_ is the average absolute abundance of ASV x in Section N of sampling sites and AA_x, OctS02_ is the absolute abundance of ASV x at OctS02, the canal head. OctS02 was excluded from Section1 for the LGF calculation. An average LGF > 0 across the three sections was considered to indicate growth in the canal. The potentially pathogenic bacterial ASVs were identified referring to a list of potentially pathogenic genera (Table S2) summarized from Woolhouse et al. (2015), and LGFs of these ASVs were examined.

## Results

### Microbiota biomass and diversity shifted with seasonality

Microbial communities showed clear seasonal shifts (Fig. 2), whether for their biomass (A and B) or alpha (C-F) and beta diversity (Fig. 2G, 2H and Fig. S1). Both the bacterial and micro-eukaryotic biomass, as measured by 16S rRNA (A) and 18S rRNA (B) gene copies, were the highest in August and the lowest in October (Fig. 2). PERMANOVA showed that seasonality significantly (*P* < 0.001) influenced both the bacterial (*F* = 22.78) and micro-eukaryotic (*F* = 25.57) community composition. Moreover, the seasonal impacts were much more pronounced than the spatial effects along the canal, whether for bacteria (*R^2^* = 0.52 vs. 0.14) or micro-eukaryotes (*R^2^* = 0.55 vs. 0.14). Accordingly, Bray Curtis-based NMDS analyses of all canal water samples (n = 52) revealed strong clustering of bacterial (Fig. 2G) and micro-eukaryotic (Fig. 2H) communities in relation to seasons, of which May and August had witnessed greater intra-season community variations along the canal. Seasonal variation was also observed for specific bacterial and micro-eukaryotic lineages (Fig. S2 and S3).

**Fig. 2.**
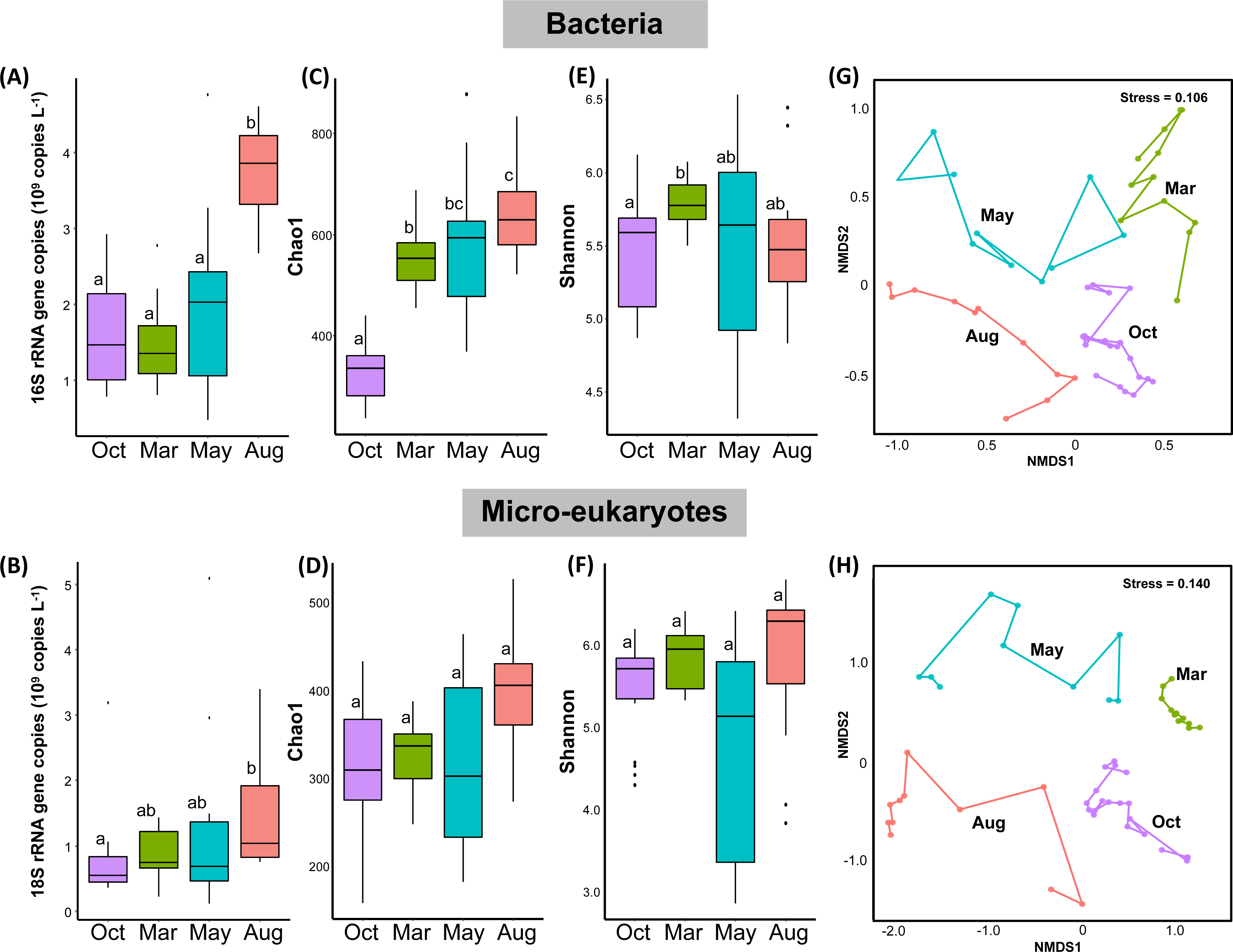
Seasonal variation of bacterial and micro-eukaryotic communities in the canal. (A-F) Ribosomal RNA gene copies (A and B), Chao1 richness (C and D) and Shannon diversity index (E and F) across seasons. Different letters above boxes indicate statistically significant differences between months of sampling (Mann-Whitney *U* test with the Benjamini-Hochberg correction, *P* < 0.05). (G-H) Ordination analysis of communities using non-metric multidimensional scaling (NMDS) based on Bray-Curtis dissimilarities. Trajectories indicate locations of sampling sites along the canal.

### Microbiota composition and diversity changed along the canal

Both the source water from Danjiangkou reservoir (S01) and the canal water were dominated by phyla *Actinobacteria* (39.7 - 65.0%, mainly hgcI clade and CL500-29 marine group), *Proteobacteria* (17.9 - 38.3%), *Bacteroidetes* (0.9 - 29.5%), *Verrucomicrobia* (1.2 - 9.2%) and *Cyanobacteria* (0.2 - 8.6%, Fig. 3A). Overall, bacterial communities at the canal head (S02) were most structurally similar to the source water (Fig. 3A and Fig. S1A). However, compared with the source water, relative abundance of *Proteobacteria* at the canal head (i.e., S02) decreased whereas *Actinobacteria* and *Bacteroidetes* increased. Within the canal, both *Cyanobacteria* (mostly *Cyanobium*_PCC-6307, *F* = 16.77, *P* < 0.001) and *Bacteroidetes* (*F* = 6.06, *P* < 0.05) showed significant decrease in relative abundance with water flow (Fig. 3A). In contrast, *Limnohabitans* (*F* = 19.87) and *Candidatus* Methylopumilus (*F* = 58.69) within *Proteobacteria* became increasingly abundant along the canal (*P* < 0.001). Likewise, both the overall bacterial biomass (Fig. S4A) and Chao1 richness (Fig. S5A) exhibited highly dynamic variation patterns along the canal. The variation trend of Shannon diversity index largely agreed with (though relatively weaker than) that of Chao1 richness (Fig. S5B).

**Fig. 3.**
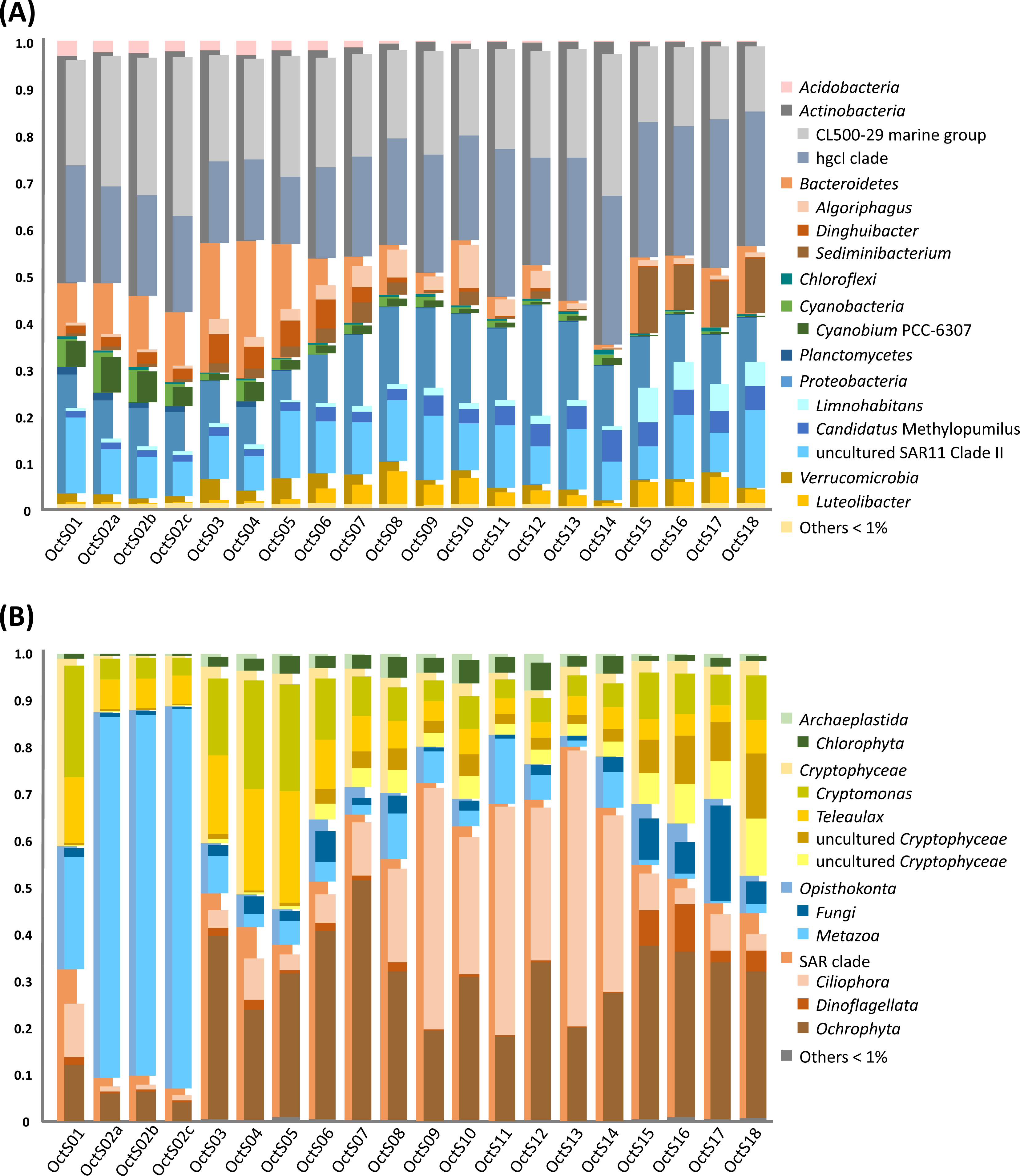
Microbial community composition and diversity along the MRP main canal (∼1432 km) of the SNWD project in October 2018. (A) Bacterial composition with abundant phyla (> 1%) and 10 most abundant genera shown. (B) Micro-eukaryotic composition with abundant phyla (> 1%) and 10 most abundant orders shown.

Unlike bacterial communities, micro-eukaryotic communities appeared more dynamic along the canal water flow (Fig. 3B). The relative abundance of *Metazoa*, mostly *Calanoida*, was as high as 24.0% in Danjiangkou Reservoir. These organisms were drastically enriched by 2.3 times upon entering the canal head site, but largely declined in the downstream sites (S03 to S18). Two predominant lineages of *Cryptophyceae*, i.e., *Cryptomonas* (*F* = 4.11, *P* = 0.06) and *Teleaulax (F* = 9.50, *P* < 0.01), were more abundant in the source water and canal upstream (S02-S08) than downstream sites (S09p-S14), accompanied by their gradual replacement by two uncultured *Cryptophyceae* lineages (*F* = 30.53, *P* < 0.001; *F* = 24.60, *P* < 0.001, respectively), especially close to the canal end (S15-S18). The canal head communities were distinct from water source and the downstream communities (Fig. 3B and Fig. S1B). This special pattern was manifested by the dominance of *Calanoida* (72.0%) at the canal head site where micro-eukaryotic biomass was much higher (3.2 ×10^9^ vs. average 5.9 × 10^8^18S rRNA gene copies L^-1^, Fig. S4B) whereas alpha diversity metrics much lower compared with the other sites (Fig. S6), suggesting intriguing local micro-eukaryotic selection in the canal head.

### Environmental factors influenced canal microbiota diversity and biomass

The influence of five environmental parameters, namely water temperature, pH, dissolved oxygen (DO), total nitrogen (TN) and fluoride (Table S3), and spatial factors (i.e., ten PCNM variables representing the quantitative spatial patterns, Table S4) on the microbial communities was tested by redundancy analysis. All the above-mentioned environmental variables and PCNM1, 2 and 4 were identified to exert significant impacts on bacterial community composition (*P* < 0.05, Fig. 4A). These variables except for PCNM4 also represented the significant influential factors for micro-eukaryotic communities (*P* < 0.05; Fig. 4B). Seasonal variation of environmental factors appeared based on the RDA plots (Fig. 4A and 4B), for instance, fluoride concentration was especially higher in March, which was supported by the significant seasonal difference in the environmental variables by Kruskal-Wallis one-way ANOVA (Table S5). The abundant actinobacterial hgcI clade (r_s_ = 0.63, *P* < 0.01) and CL500-29 (r_s_ = -0.68, *P* < 0.01) marine group were among the bacterial lineages positively and negatively related to fluoride concentration, respectively (Table S6). For micro-eukaryotes, for instance, an ASV affiliated to *Mediophyceae* (r_s_ = 0.66, *P* < 0.01) was positively correlated with fluoride (Table S7).

**Fig. 4.**
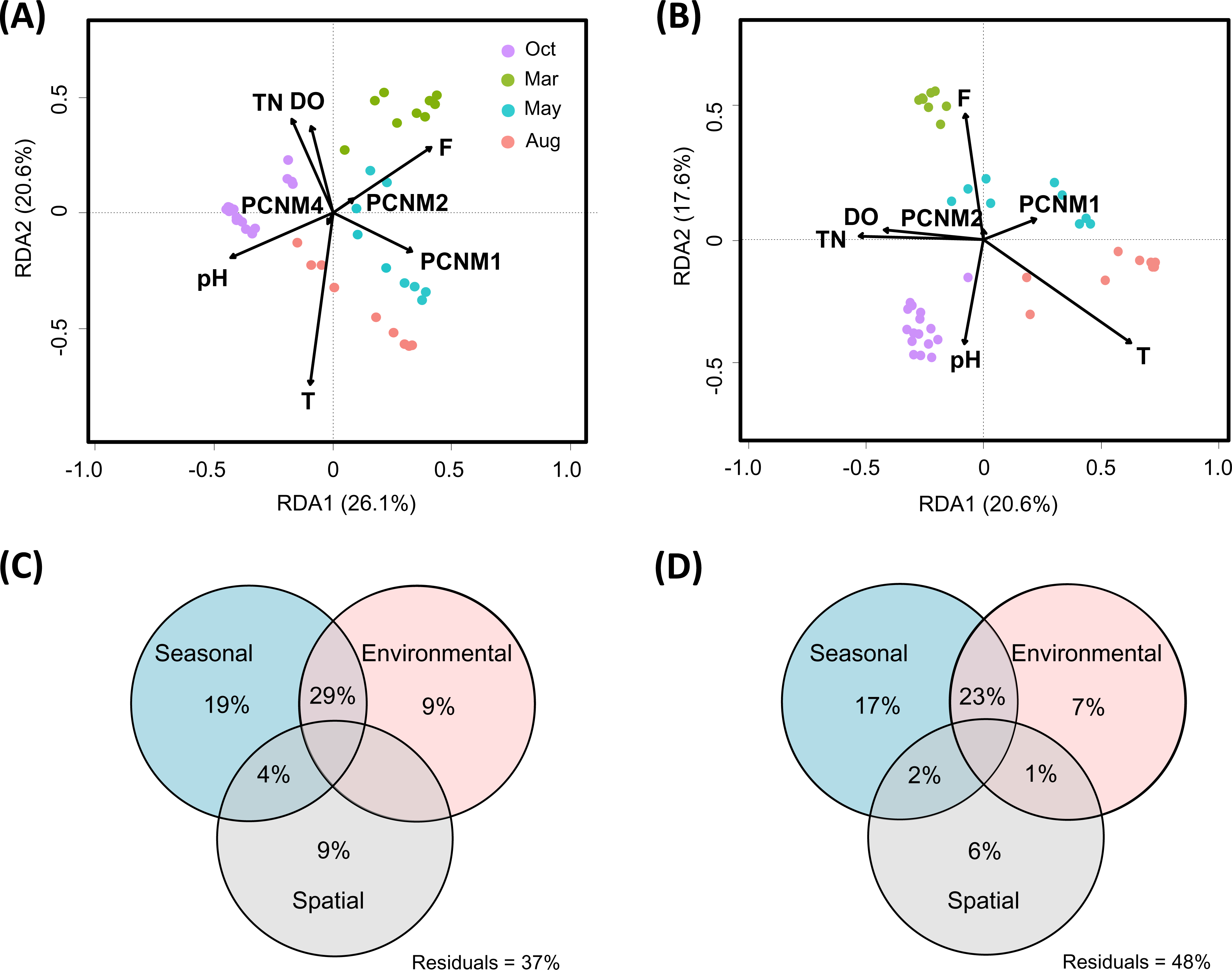
Redundancy analysis (RDA) and variation partitioning analysis (VPA) demonstrating the relative influence of seasonality, spatial and environmental factors on the canal microbial communities. The RDA biplots show factors impacting bacterial (A) and micro-eukaryotic (B) communities. The VPA diagrams show the percentage contribution of seasonal, spatial and environmental factors to bacterial (C) and micro-eukaryotic (D) community variations. Only those factors identified to significantly influence bacterial and micro-eukaryotic communities are shown in the RDA and included in the VPA analyses. PCNMs, geographic factors generated by principal coordinate analysis of neighborhood matrices. T, temperature; F, fluoride; TN, total nitrogen; DO, dissolved oxygen.

The identified significant variables were employed in further variation partitioning analysis (VPA). The result showed clearly stronger influence of seasonality than that of spatial and examined environmental factors for both bacterial and eukaryotic communities (Figure 4C and 4D). Seasonality alone could explain 19% and 17% of the bacterial and micro-eukaryotic community variance, respectively, while only 6 - 9% of the variance could be explained exclusively by environmental or spatial factors. Notably high proportions of the community variance could be attributed to the co-effects of seasonality and environmental factors (29% for bacteria and 23% for micro-eukaryotes), implying their strong interplay in shaping microbial communities. Besides, temperature, DO and TN were found to be significantly correlated with bacterial biomass, while only DO for micro-eukaryotic biomass (as measured by 16S rRNA and 18S rRNA gene copies, Table S8).

### Bacterial and micro-eukaryotic communities exhibited distinct and spatio-temporally dynamic stochasticity

The apparent dispersal of microorganisms in the canal water and engineered hydraulic manipulations (i.e., inverted siphon, aqueduct, and tunnels) along the main canal drove us to question on the relative contribution of stochastic vs. deterministic processes to the canal bacterial and micro-eukaryotic community assembly, which was first quantitatively assessed with a recently proposed metric, normalized stochasticity ratio (NST) (Ning et al., 2019). This new metric quantifies the relative importance of stochasticity in governing community assembly ranging from 0 to 1, with a value > 0.5 indicating more stochastic assembly and < 0.5 more deterministic assembly. NST analysis revealed contrasting trends between bacteria and micro-eukaryotes. In the upstream section, a highest NST of 0.78 was estimated for bacteria (Fig. 5A). This value decreased to 0.46 in the middle section, and to as low as 0.32 close to the canal end. On the contrary, NST was generally above 0.5 for micro-eukaryotes throughout the canal, with the highest value of 0.75 in the middle section (Fig. 5A).

**Fig. 5.**
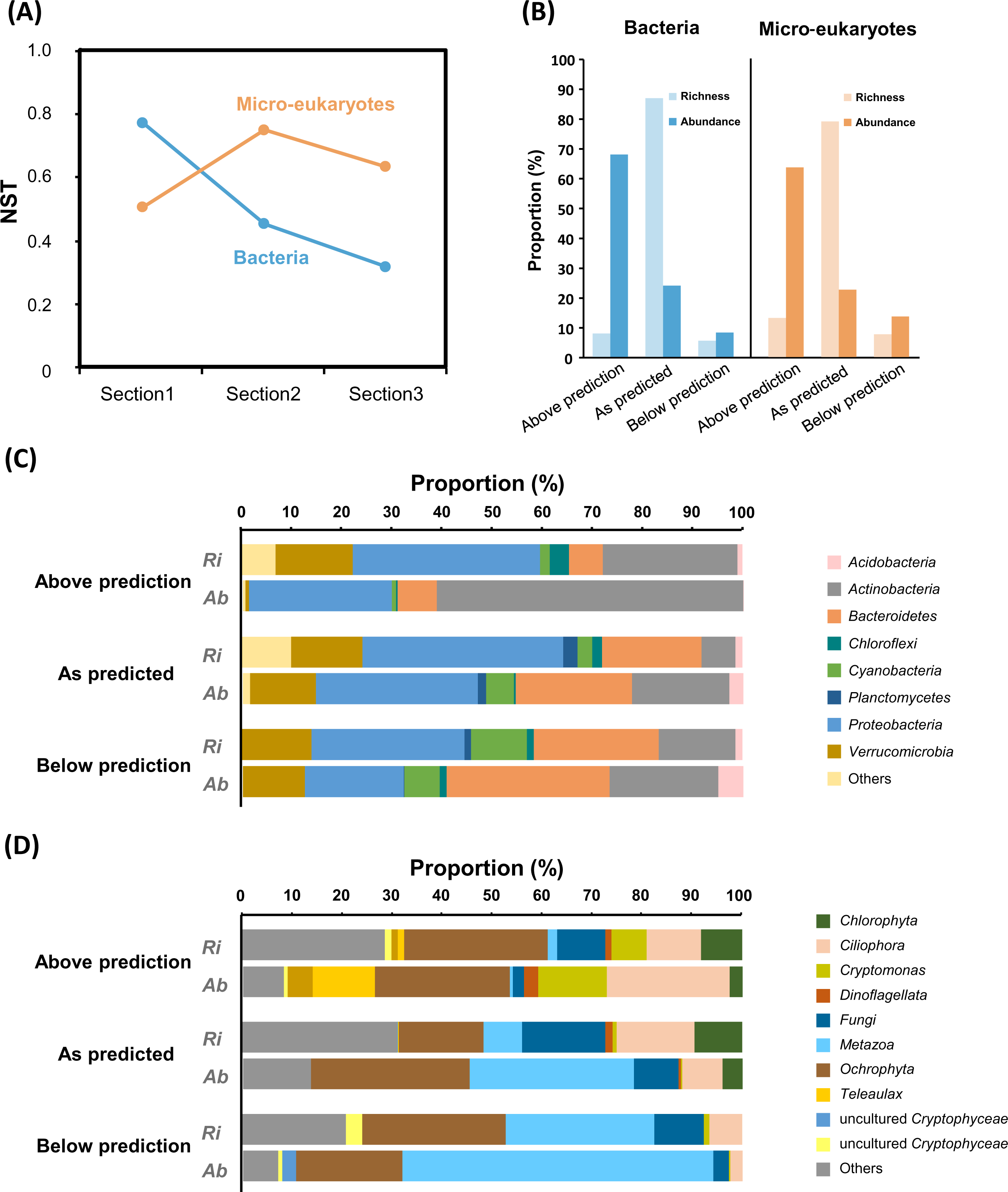
Normalized stochasticity ratio (NST) estimation and microbiota partitioning by population compliance to the neutral model prediction. (A) NST estimated stochasticity in bacterial and micro-eukaryotic community assembly. (B) Proportions in richness (ASV number) and abundance (sequence number) of the three fitting groups (occurrence frequency above prediction, as predicted and below prediction) within the microbial communities to the Sloan neutral model. (C and D) Taxonomic composition of the three fitting partitions in richness (Ri) and abundance (Ab) for bacterial (C) and micro-eukaryotic (D) communities. Only abundant bacterial phyla (> 1%) and 10 most abundant micro-eukaryotic orders are shown.

While NST provides rigorous quantification of relative importance of stochasticity, the Sloan neutral model analysis (Sloan et al., 2006) allows for assessing how well the observed community patterns could be explained by neutral processes (e.g., birth, death, and immigration) and attributing the observed community-level assembly pattern to the divergent population behavior. According to seasonal comparison of the model fitness, neutral model could explain much higher proportions of community patterns in October (R^2^= 0.73 and 0.67 for bacteria and micro-eukaryotes, respectively) and March (R^2^ = 0.77 and 0.72), but lowest in May (R^2^= 0.42 and 0.13, Fig. S7).

To link community patterns to the underlying population dynamics, bacterial and micro-eukaryotic ASVs were classified into three groups based on the neutral model analysis of the October samples. Depending on whether they empirically occurred more frequently than, as frequently as or less than predicted by the neutral model, ASVs were assigned to “above prediction”, “as predicted” and “below prediction” groups that were different in abundance and composition (Fig. 5B-D). We found that 7.8% of the bacterial community richness (i.e., number of ASVs) accounted by above-prediction group corresponded to 67.9% of the total number of sequences, which was dominated by *Actinobacteria* and *Proteobacteria*, whereas the neutrally distributed group contributed 86.8% of the richness but only 23.9% of the abundance (Fig. 5B and Fig. S8). Similar trend was detected for micro-eukaryotes (Fig. 5B). The three partitioning groups of ASVs also showed distinct taxonomic composition (Fig. 5C, 5D and Fig. S8). *Cryptomonas* constituted 13.7% of the above-prediction group in abundance, but they were almost absent in the neutral and below-prediction groups. *Metazoa*, in contrast, accounted for only 0.6% of the above-prediction group, but as high as 32.9% and 62.3% of the abundance of the neutral and below-prediction groups (Fig. 5D).

### Selective growth of immigrating bacterial and micro-eukaryotic populations along the canal

To resolve which populations of the water communities died off, sustained, or proliferated during dispersal over 1432-km canal, we proposed local growth factor (LGF) (see equation 1 in Methods) as a quantitative measure of variation in population abundance in each canal section compared to the abundance at the canal head, with an average LGF > 0 across the three sections indicating growth in the canal. LGF analysis was applied with the Oct sample dataset composed of 1389 bacterial and 1229 eukaryotic ASVs, using absolute abundance of ASVs estimated by multiplying relative abundance of ASVs at each canal site with the total copies of bacterial 16S rRNA (7.89 ×10^8^to 2.93×10^9^ copies L^-1^) or micro-eukaryotic 18S rRNA (3.62×10^8^ to 3.19 ×10^9^ copies L^-1^) genes therein. Micro-eukaryotes showed higher overall occurrence (0.313±0.008 vs. 0.216±0.007; *P* < 0.001) while lower variability (2.879±0.033 vs. 3.055±0.032; *P* < 0.001) in absolute abundance compared to bacteria.

In total, 67.1% of the bacterial ASVs (Table S9) and 82.3% of the micro-eukaryotic ASVs (Table S10) displayed positive average LGFs. The ASVs were further observed to form clearly three clusters based on their LGFs in each section: LGF > 3, -3 < LGF < 3 and LGF < -3, which could indicate substantial growth, survival, and die-off of the microbial populations (Fig. 6). The survival group represented the largest proportion of the ASVs (45.6 - 60.2%), while only 4.1 - 21.7% of the ASVs went extinction in the canal. We also found that both the contrastingly high and low LGFs tended to be attributed to high-abundance microbial lineages (Fig. 6A and B). For example, the abundant *Luteolibacter* sp. (ASV106) and *Limnohabitans* sp. (ASV27) exhibited the highest LGF values in Section 2 and Section 3, respectively (Table S9). In stark contrast, ASV138 assigned to actinobacterial CL500-29 marine group and two *Cyanobium* PCC-6307 lineages (ASV144 and ASV250) were especially restrained in growth close to the canal end (Table S9). A total of 53 bacterial ASVs in the canal were related to 16 genera included in a published reference pathogen list (Table S2, summarized from Woolhouse et al., 2015). While LGF of most (39 out of 53) of these potentially pathogenic ASVs fell between -3 and 3, three *Acinetobacter*-associated ASVs (ASV794, ASV1108 and ASV 1261) showed LGFs ranging from -5.9 to -6.1, indicating notable depletion along the canal. For micro-eukaryotes, two uncultured *Cryptophyceae* lineages (ASV54 and ASV80) as well as two ASVs related to *Acineta flava* (ASV224) and *Spumella* sp. (ASV59) were elevated throughout the canal (Table S10).

**Fig. 6.**
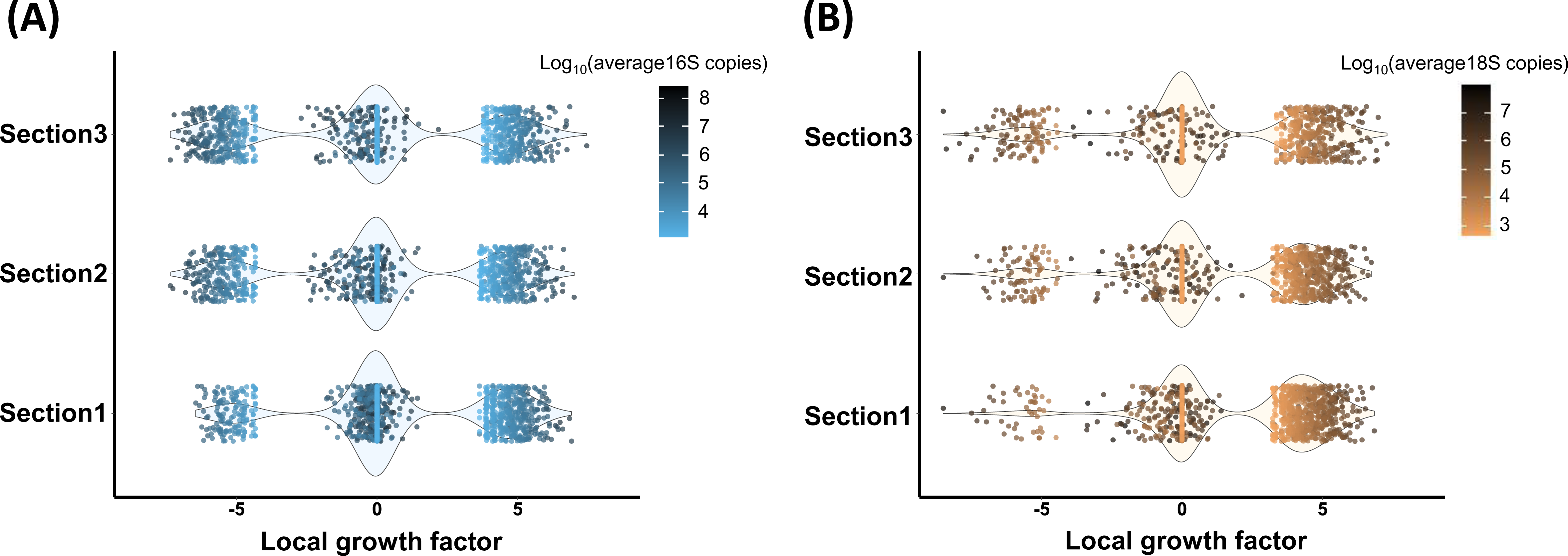
Microbial population proliferation across canal sections evaluated by local growth factor analysis. Distribution patterns of population local growth factors in the different canal sections are shown with violin plots for bacteria (A) and micro-eukaryotes (B). Each dot represents an ASV with colors indicating its average abundance across the canal.

## Discussion

Diversion canals widely built for reallocation of water resources represent an ideal engineered system to model microbial communities fulfilling key eco-environmental services, as well as reflecting changes in ecosystem structure and function. Based on quantitative metagenomic approaches that enable absolute quantification of microbial taxon, we uncovered the microbiota spatio-temporal dynamics, key processes driving community assembly, and navigation of immigrating microorganisms through the world’s largest water diversion project.

### Microbiota composition and biomass are dynamic along the canal

Both bacterial and micro-eukaryotic communities in the source water showed marked variations in taxonomic composition upon entering the canal, which could be attributed to the readily comprehensible differences in the environmental conditions between the natural reservoir and its receiving engineered canal and potential influences exerted by the approach channel through which source water enters the canal. Meanwhile, we observed intriguingly decreased water microbial diversity after water transportation through an inverted siphon (S09 vs. S09p, Fig. S5 and S6). Taken together, these suggest that hydraulic structures along the canal apply considerable influence on the microbial community diversity and composition, which needs to be fully considered when setting baseline communities for long-term microbial monitoring in the water diversion canal.

The prevalence of uncultured actinobacterial hgcI clade and CL500-29 marine group echoed what was previously reported in other freshwater habitats (Keshri et al., 2018; Yang et al., 2017). While both groups of yet-to-be-cultured lineages have been shown to favor lower level of dissolved organic carbon and temperature (Lindh et al., 2015), the hgcI clade was additionally implicated to be competitively advantageous in utilizing low-concentration organic carbon (Glöckner et al., 2000) and to harbor potentially photoheterotrophic species showing preferential utilization of nitrogen-rich organic compounds in freshwater (Ghylin et al., 2014), implying its potential role in the nutrient uptake and water auto-purification.

Unlike bacterial communities, micro-eukaryotic communities in the canal have not yet been explored before. By virtue of the high-throughput molecular analyses, we detected not only *Rotifea*, copepods and protozoa previously microscopically observed as abundant in the Danjiangkou Reservoir (Li et al., 2012), but also thousands of micro-eukaryotic populations (as ASVs), of which 6.7% are uncultured or unclassified even at the class level, implying detection of numerous novel populations in the canal water. Notably, metazoan *Calanoida* was repeatedly found to show drastic increase in abundance in the micro-eukaryotic communities of canal head, compared with in the source water in both October and August. This big order of aquatic crustaceans belonging to *Copepoda* inhabiting marine and freshwater environments are found as the most successful among zooplankton colonizing pelagic habitats (Blanco-Bercial et al., 2011; Bradford-Grieve, 2002). We further noted that the SAR clade and *Cryptophyceae* showed substantially decreased relative abundance as a result of high enrichment of *Calanoida* compared with the source water. However, their absolute abundance or biomass, as we estimated as their 16S rRNA gene copies, remained comparable (2.78 vs. 3.4 ×10^8^16S-copies L^-1^for the SAR clade, and 3.68 vs. 4.25×10 16S-copies L^-1^for *Cryptophyceae*). This strong decoupling pattern between relative and absolute abundance cannot have been resolved by currently popular sequence-based relative measures alone which have been shown to provide evidently misleading information on the variation of absolute abundance (Stämmler et al., 2016; Vieira-Silva et al., 2019). It also well demonstrates the power of our quantitative metagenomics approaches as an alternative to a more direct but difficult-to-achieve quantitative PCR assay approach (due to the lack of 18S rRNA gene universal primers) for estimating total micro-eukaryotic biomass.

### Microbiota assembly is deterministically driven by seasonally dependent habitat filtering effects

Elucidating mechanisms and factors shaping biogeographic patterns has become a central goal in microbial ecology (Hanson et al., 2012). Environment exerts influence on community composition by selecting for populations that better adapted to the corresponding ecological niches. We found that temperature, pH, dissolved oxygen, total nitrogen and fluoride all showed significant effects on the community dynamics in the main canal. Fluoride affects bacterial growth and metabolic activity through targeting enzyme inhibition (Liao et al., 2015). Fluoride has been reported to alter microbial community in high-fluoride groundwater (Zhang et al., 2019a) and to exhibit inhibitory effect on bacterial populations involved in various wastewater treatment processes (Ochoa-Herrera et al., 2009). Our results showed that fluoride with concentration falling within the range of 0.14-0.22 mg L^-1^(Table S3) could also affect microbial community structure in a drinking water diversion system, where fluoride could be mainly contributed by natural processes, such as mineral weathering (Nong et al., 2019), and atmospheric pollutants (Pu et al., 2017). Interestingly, we found positive response of the hgcI clade to elevated fluoride concentration (Table S6). While the mechanisms of their fluoride resistance still await exploration, such evidence is in line with their favorable distribution in a wide range of water ecosystems.

Although bacterial and micro-eukaryotic communities are both significantly influenced by the examined environmental variables and spatial segregation, their overall variation were explained more by seasonality. Meanwhile, the strong co-influence of temporal and environmental factors revealed by variation partitioning analysis could be at least partially explained by seasonal variation of the examined variables (Table S5), suggesting season-driven habitat filtering effects on microbiota. These results together imply a strong interplay between temporal, spatial and environmental factors on influencing microbial communities in the canal water, as previously noticed in natural lakes (Hu et al., 2017). Although potential contribution from unmeasured environmental factors cannot be ruled out, the remaining unexplained variation could possibly be generalized and linked to the influence of stochastic processes.

### Contrasting proportion of stochasticity contributes to bacterial and micro-eukaryotic community assembly

Microbial community assembly is typically governed by four processes: selection, drift, speciation and dispersal (Vellend and Agrawal, 2010). While selection is inherently deterministic and drift is stochastic, speciation and dispersal could involve both deterministic and stochastic mechanisms (Chase and Myers, 2011). For the MRP main canal, an investigation period of one year is probably short for resolving speciation. Thus, selection (deterministic processes), drift (stochastic processes) and dispersal (involving both) are expected to shape the observed community assembly patterns, and their relative contributions could be disentangled to help answer when, where, why and how much each process contribute to the observed patterns (Hanson et al., 2012).

Based on NST estimation and neutral modeling, we found that relative importance of stochastic and deterministic processes in microbial community assembly differ between bacteria and micro-eukaryotes, and among seasons and the canal sections. Neutral processes explained ∼67% to 76% of the community patterns in October and March, but notably less in May (∼41% and 13% for bacteria and eukaryotes, respectively). Liu et al. (2018a) reported a prominent shortage in water storage of Danjiangkou Reservoir related to seasonal flood control in May, which could have possibly imposed higher environmental selection effect to the microbiota in the reservoir and across the canal, making the observed decreased community stochasticity (or increased determinism) plausible. Although ecological stochasticity adds to challenge to predict microbial communities, the new knowledge produced in this study undoubtedly contribute to an improved understanding on when the canal water microbiota is more prone to stochastic processes, or under which conditions stochasticity is of more relevance.

The NST values suggested that the relative contribution of deterministic processes is larger to bacterial community assembly compared with micro-eukaryotic communities, where stochasticity prevails. Recently, Gweon et al. (2021) reported that stochastic processes played a more important role in the planktonic bacterial communities in the River Thames, presumably due to the influence of various natural and anthropogenic sources of the surface water. What we found showed that in a water ecosystem almost isolated from other sources than resource water, bacterial communities are predominantly shaped by deterministic abiotic and biotic processes. The micro-eukaryotes, however, possess structurally and functionally more complex cells than prokaryotes (Massana and Logares, 2013), which could facilitate niche adaptation and diminish the environmental filtering effect at the community level. This was supported by our observation that the micro-eukaryotes were distributed with higher occurrence frequency and lower variability across the canal compared to bacteria. The different controlling mechanisms between bacterial and micro-eukaryotic community assembly found here has been observed in other natural habitats, such as Antarctic (Logares et al., 2018) and Tibetan lakes (Liu et al., 2020b). Our comparative analyses between bacteria and micro-eukaryotes across canal sections and over seasons thus provide first informative insights into the domain-specific, and space- and season-dependent patterns in the canal water community assembly.

### Divergent population lifestyles underly microbiota dynamics revealed by absolute quantification

A synthesis of population and community ecological perspectives can provide valuable insights into how microorganisms drive ecosystem processes (Koch et al., 2018). Microbial populations in this giant system displayed interestingly divergent lifestyles reflecting the difference in their adaptability to the canal environment. The Sloan neutral model sorted the microbial ASVs to above-prediction, below-prediction (jointly non-neutral) and as-predicted (neutral) groups, which were different in community composition (Fig. 5C and 5D), indicating taxonomically dependent competitive fitness in the investigated canal. In addition, we observed contrasting low (∼20%) contribution of neutral group to community abundance despite their high proportion in richness (∼80%) for both bacteria and micro-eukaryotes (Fig. 5B). Chen et al. (2019) reported for micro-eukaryotic communities in a subtropical river as source of drinking water the dominance of neutral groups in both richness and abundance of the overall communities as well as across taxonomic groups. These contrasting patterns suggested that microorganisms are prone to more selective processes in an engineered canal compared with natural habitats.

In particular, we demonstrate the use of local growth factor (LGF) as a meaningful metric to infer growth of immigrating microbial taxa in the canal by employing absolute quantification of microbial populations. This leads to the first estimation that 67.1% of bacterial and 82.3% of micro-eukaryotic populations were capable of sustaining growth (LGF > 0), implying again better acclimation of eukaryotes to the environmental conditions in the canal. It is worth clarifying that the microbial growth estimated here was aimed to characterize fate of immigrating microorganisms across the canal, calculated based on relative fluctuation in absolute abundance at local sites compared with canal head, and thus is not equivalent to per-capita or population growth in the Sloan neutral model (Sloan et al., 2006). *Limnohabitans* sp., ubiquitous in a wide variety of freshwater habitats, showed remarkable growth across the canal. Members of *Limnohabitans* typically display high growth rate as well as metabolic flexibility, and they actively utilize alga-derived dissolved organic substances and are vulnerable to protist grazing (Horňák et al., 2017; Šimek et al., 2013; Šimek et al., 2011). *Cryptophyceae* were among the micro-eukaryotes that displayed prominent growth throughout the canal, which are small flagellates as important food of many protists and micro-crustaceans (Horňák et al., 2017; Posch et al., 2015), and are perceived as characteristic of clean water (Lackey, 1941). Thus, *Limnohabitans* and *Cryptophyceae* are potentially important constituents of water food webs in the investigated canal. Two cyanobacterial lineages (*Cyanobum* PCC-6307) and *Acinetobacter* spp., on the contrary, were clearly depressed in the canal downstream sections, demonstrating the elimination of these microorganisms of water security concern along the canal. *Acinetobacter* spp. are ubiquitous in nature of major ecological and clinical significance, featuring, for instance, multi-antimicrobial resistance and rapid resistance development (Jung and Park, 2015). The diminishing of the versatile *Acinetobacter* spp. along the canal could possibly be attributed to unfavorable environmental conditions (Al Atrouni et al., 2016) or composite intra- and inter-domain biological interactions. Further research efforts with genome-centric metagenomic approaches are needed to elaborate the mechanisms behind such die-off patterns. It is noteworthy that due to the restriction in taxonomic resolution of the ribosomal RNA gene sequencing analysis employed in this study, species-level confirmation is still needed to evaluate the relevance of these potentially pathogenic microorganisms. Together, our results demonstrated the power of local growth factor to identify key players in the investigated system and facilitate predicting fate of the microbes traversing such thousand-kilometer water diversion canal.

## Conclusions

The main canal of the Middle Route Project of the South-to-North Water Diversion projects is an engineered system where the aquatic ecosystem is still at its early forming stage (since 2014). Various aspects of hydrology, geochemistry and ecology need to be intricately elaborated to jointly contribute to sustained system’s eco-environmental services, especially water auto-purification ensuring its quality and competence for designed social-economic aims. The present study uncovered the underlying microbial community assembly mechanisms and population dynamics in a giant engineered water diversion water canal, such as the sharp decline of potentially harm-producing microbes (e.g., *Cyanobacteria*) and potential pathogens (e.g., *Acinetobacter* spp.), and the favorable enrichment of alga-derived organic scavengers (e.g., *Limnohabitans*) as potential key microbial food-web players, demonstrating the valuable insights provided by microbial community and population assessments for water monitoring and their potential in indicating ecosystem disturbance. Furthermore, the bacterial community shift driven by combined environmental effects as well as the stochasticity in micro-eukaryotic community assembly needs to be considered in water quality monitoring along the canal, towards an appropriate microbe-based biomonitoring which enables deeper understanding of processes behind water quality parameters. Considering possibly occasional pollution into the canal water caused by, for example, traffic accidents on highway bridges across the canal, and the central roles microorganisms play in the water quality maintenance, future efforts are needed to explore the metabolic functions of microbial community contributing to water purification (e.g., carbon, nitrogen, and phosphorus removal) during diversion and sustaining water quality.

## Acknowledgments

We would like to thank Yuanzhen He, Guoqing Zhang, Zhipeng Zhang, Shengguang Yuan, Aoxiang Fan and Qinglin Jia for the assistance in water sampling. This work was supported by National Natural Science Foundation of China (Grant No. 51908467 and No. U2040210), and Zhejiang Provincial Natural Science Foundation of China (Grant No. LQ20C030002).

