## Supplementary Methods, Figures and Tables for "Untangling Microbiota Diversity and Assembly Patterns in the World’s Largest Water Diversion Canal"

**Method S1** Study sites, sampling and physiochemical monitoring

**Method S2** DNA extraction, PCR amplification and sequencing

**Method S3** Bioinformatics and statistical analyses of microbial diversity

**Method S4** Neutral and null model analyses of microbial community patterns

**Method S5** Population lifestyle and local growth factor analyses

**Figure and Table Legends**

**Fig. S1** Bray-Curtis similarity-based dendrogram showing bacterial (A) and micro-eukaryotic (B) community composition for all the four sampling campaigns. Sample names are colored according to month of sampling. Relative abundance of the top 10 abundant phyla for bacteria and all the phyla for eukaryotes were shown in bar plots.

**Fig. S2** The distribution of top 30 abundant bacterial genera across the canal.

**Fig. S3** The distribution of top 30 abundant micro-eukaryotic order-level lineages across the canal.

**Fig. S4** Bacterial 16S (A) and eukaryotic 18S (B) rRNA gene copies quantified for microbial communities along the canal across the four sampling campaigns.

**Fig. S5** The Chao1 and Shannon indices of bacterial communities along the canal across the four samplings.

**Fig. S6** The Chao1 richness and Shannon diversity index of micro-eukaryotic communities along the canal across four samplings.

**Fig. S7** Fit of the neutral model to Oct (A, B), Mar (C, D), May (E, F) and Aug (G, H) samples across the canal for bacterial and micro-eukaryotic communities. The solid blue lines indicate the prediction of ASV occurrence frequencies predicted by the model, and the dashed blue lines represent 95% confidence intervals. ASVs that occur more frequently, less frequently than predicted or as predicted by the model are shown in different colors. Parameter “m” represents estimated migration rate.

**Fig. S8** Contrasting taxa-resolved proportions of the three fitting partitions (occurrence frequency above prediction, as predicted and below prediction by the Sloan neutral model) in richness (ASV number) and abundance (sequence number) for bacterial and micro-eukaryotic communities. Only abundant bacterial phyla (> 1%) and 10 most abundant micro-eukaryotic orders were shown.

**Table S1** Ratio of bacterial 16S and eukaryotic 18S rRNA gene copies along the canal based on metagenomic sequencing of the canal water samples.

**Table S2** List of potentially pathogenic genera summarized according to Woolhouse et al. (2015). Genera highlighted in red were detected in this study.

**Table S3** Physiochemical properties of water samples collected from 16 sites over four seasons.

**Table S4** PCNM (Principal Coordinates of Neighborhood Matrices) of the 16 sites where water properties were measured.

**Table S5** Seasonal variation of environmental variables. Significance of seasonal effects were tested with the Kruskal-Wallis one-way ANOVA.

**Table S6** Bacterial ASVs with significant and strong correlation with fluoride. Correlation was tested by computing Spearman’s rank correlation, corrected with the Benjamini-Hochberg method. Only ASVs with Spearman’s r_s_ > 0.6 or < -0.6, and *P* < 0.05 are shown.

**Table S7** Micro-eukaryotic ASVs with significant and strong correlation with fluoride. Correlation was tested by computing Spearman’s rank correlation, corrected with the Benjamini-Hochberg method. Only ASVs with Spearman’s r_s_ > 0.6 or < -0.6, and *P* < 0.05 are shown.

**Table S8** Correlation between ribosomal RNA gene copies, diversity metrics and environmental variables.

### Supplementary Methods

#### Method S1 Study sites, sampling and physiochemical monitoring

The Middle Route Project (MRP) of the South-to-North Water Diversion (SNWD) Projects started from Danjiangkou reservoir and diverted water northwards until Beijing and Tianjin. Started to operate in 2014, the MRP is an artificial system without direct connection with surface waters in the area. A total of 50 samples were collected from 19 sites (Fig. 1), including one at the Danjiangkou reservoir (S01) and 18 along the main canal of the MRP (S02-S18), among which 18 sites were sampled during October to November 2018 (Oct), 11 in March (Mar), 10 in May and 11 in August (Aug) 2019. Higher chances of algal bloom were suggested in the source reservoir in summer and autumn (May and Aug; Shen et al., 2015), while Mar and Oct were viewed as to represent time before and after these periods. The S09p and S09 represent two adjacent sampling sites in the upstream and downstream of an inverted siphon, and an aqueduct is located between S11 and S12. The 18 sampling sites along the canal were grouped to three sections: S02-S08 (Section1), S09p-S14 (Section2) and S15-S18 (Section3). Canal water (~1 L) was sampled and filtered through 0.22-μm cellulose nitrate filters (Sartorius, Germany) before storage at -20 °C until further operation. Physiochemical properties, including water temperature (T), pH, dissolved oxygen (DO), total nitrogen (TN) and fluoride (F^-^), were measured according to the environmental quality standard for surface water of China (GB3838-2002). Briefly, T, pH and DO were measured *in situ* using Hydrolab DataSonde 5. TN was determined via alkaline persulfate digestion followed by UV spectrophotometry. The concentrations of F^-^ was measured by ion chromatography.

#### Method S2 DNA extraction, PCR amplification and sequencing

Genomic DNA was extracted from duplicate filters per sample using FastDNA Spin Kit for Soil (MP Biomedicals, USA) following the manufacturer’ s protocol. Except that triplicate DNA extracts were sequenced for OctS02 (OctS02a, b and c), duplicate DNA extracts per sample were pooled for sequencing. Bacterial 16S rRNA gene V3-V4 region and eukaryotic 18S rRNA gene V4 region were amplified using primer pairs 338F (5’-ACTCCTACGGGAGGCAGCA -3’), 806R (5’-GGACTACHVGGGTWTCTAAT-3’) and 528F (5’-GCGGTAATTCCAGCTCCAA-3’), 706R (5’-AATCCRAGAATTTCACCTCT-3’) with barcode, respectively. The pooled amplicon library was sequenced on the Illumina Hiseq 2500 platform for Oct samples and Novaseq platform for Mar, May and Aug samples, producing 250 bp paired-end reads at the Guangdong Magigene Biotechnology Co., Ltd. (Guangzhou, China).

Real-time quantitative PCR was performed to quantify the bacterial biomass in each genomic DNA sample, as described in Ju et al. (2019). Each DNA extract was assessed in two dilutions with at least duplicate measurements on a qTOWER3 thermal cycler (Analytik Jena, Germany). The primers used for 16S rRNA gene-targeted qPCR were Ba349F (5’-AGGCAGCAGTDRGGAAT-3’) and Ba806R (5’-GGACTACYVGGGTATCTAAT-3’), with Taqman probe Ba516F (5’-FAM-TGCCAGCAGCCGCGGTAATACRDAG-TAMRA-3’)(Takai and Horikoshi, 2000). DNA template (2 μl) was quantified in each 25-μl reaction with Premix Ex Taq (Probe qPCR; Takara, Dalian, China) following the manufacturer’ s instructions. Plasmid with the target fragment covering the primer sites was used as standard, in a concentration series between 10^7^ and 10^0^ copies μl^-1^.

Each genomic DNA was also used for shot-gun metagenomic sequencing. Libraries were generated from 1 μg DNA per sample using NEBNext® Ultra™ DNA Library Prep Kit for Illumina (NEB, USA), following manufacturer’s recommendations. The constructed libraries were sequenced on the Illumina’s Novaseq platform using a 150 bp paired-end sequencing strategy at the Novogene (Beijing, China). The sequencing data generated in this study have been deposited into CNGB Sequence Archive (CNSA) (Guo et al., 2020) of China National GeneBank DataBase (CNGBdb) (Chen et al., 2020) with accession number CNP0001593.

#### Method S3 Bioinformatics and statistical analyses of microbial diversity

The 16S and 18S rRNA gene amplicon sequence data were processed using Qiime2 pipeline (Bolyen et al., 2019) and DADA2 algorithm (Callahan et al., 2016). After quality trimming, merging of paired sequences and removal of chimeric sequences, amplicon sequence variant (ASV) tables were generated based on 100% sequence similarity for 16S rRNA and 18S rRNA genes. Taxonomic assignment of the ASVs was conducted using the Silva database (Quast et al., 2013) (version 132). The ASVs annotated as chloroplasts or mitochondria were excluded from further analyses. The ASV tables were then rarefied to an even sequence depth of 31727 and 38888 for the 16S and 18S data, respectively. The final ASV tables consisted of 6705 bacterial and 3217 eukaryotic ASVs, respectively.

The alpha-diversity metrics (i.e., Chao1 richness and Shannon diversity index) of bacterial and micro-eukaryotic communities were computed in Qiime2. All the following univariate and multivariate statistics were performed in R (R Core Team, 2020). Pairwise seasonal difference in ribosomal RNA gene copies, Chao1 richness and Shannon diversity index was evaluated with Mann-Whitney *U* test, with *P*-values corrected for multiple testing with the Benjamini-Hochberg method. Correlation between microbial biomass (as 16S and 18S copies), alpha-diversity metrics and environmental variables was tested by computing Spearman’s rank correlation. Spearman’s rank correlation was also calculated and tested between microbial ASVs and environmental variables, corrected with the Benjamini-Hochberg method. Seasonal effects on the environmental variables were tested with the Kruskal-Wallis one-way ANOVA. Linear regression model was applied to test the increase or decrease in relative abundance of microbial lineages along the canal. Non-metric multidimensional scaling (NMDS) was performed to compare the composition of bacterial and eukaryotic communities along the canal and across the seasons based on Bray-Curtis similarity. Bray-Curtis similarity was also used to hierarchically cluster the communities, generating dendrograms for visualization. Significant clustering of the microbial communities by geographic location and impact of sampling season was tested using PERMANOVA with 999 permutations. The significance of individual environmental and spatial variables in explaining community variation was evaluated by redundancy analysis (RDA) based on Hellinger-transformed relative abundance matrices of bacterial and micro-eukaryotic ASVs. A set of spatial factors was generated for the sampling sites based on the geographic coordinates using the principal coordinates of neighboring matrices (PCNM) analysis (Borcard and Legendre, 2002). Spatial and environmental factors found to be significant in explaining community variation in the RDA were included for variation partitioning analysis (VPA), applied to compare the impact of temporal, spatial and environmental factors on microbial communities.

Quality filtered metagenomic data were ublast (Edgar, 2010) against the Silva database (Quast et al., 2013) (version 132) to identify 16S rRNA and 18S rRNA gene reads. The 16S rRNA and 18S rRNA gene copies in each metagenome was estimated as the sum for the read coverage over the full length of each matched reference sequence in the database using the following equation:

$Copy numbers=\sum_{1}^{n} \frac{L_{\mathrm{Reads}}\times N}{L_{\mathrm{Genes}}}$	(1)

where L_Genes_ is the length of a 16S rRNA or a 18S rRNA gene reference sequence in the SILVA database; L_reads_ is sequencing length of metagenomic reads; N is the number of sequencing reads in a sample with one specific 16S rRNA or 18S rRNA gene reference sequence as the best hit; n is the number of 16S rRNA or 18S rRNA gene reference sequences in the database. The copies of 18S rRNA genes in each water sample were estimated as 16S rRNA gene copies (determined by the aforementioned qPCR analysis) divided by ratios of 16S rRNA to 18S rRNA gene copy (Table S1) as sequenced and estimated in the metagenomic data.

#### Method S4 Neutral and null model analyses of microbial community pattern

To disentangle the relative contribution of stochastic and deterministic processes to microbial community assembly, Sloan neutral model and null model-based NST approach were applied to the 16S and 18S community data. The Sloan Neutral Community Model (Sloan et al., 2006) describes how microbial communities can be explained by stochastic (birth-death immigration) processes. This model assumes that individual populations with higher average abundance across all samples should be more frequently observed. The Sloan neutral model was fit to the observed occurrence frequency of ASVs and their mean abundance across communities (representing abundance in the metacommunity) with a parameter *m* (migration rate) describing the probability that a dead individual is instantly replaced by an immigrant from the metacommunity rather than by reproduction of a member from the local community. Fitting of the model and calculation of 95% confidence intervals around the model predictions were performed in R according to Burns et al. (2016). Neutral model analysis was applied to both bacterial and eukaryotic ASV datasets across the canal and in different canal sections. Accordingly, ASVs from each dataset were classified into three groups depending on whether their observed occurrence falling higher than (“above prediction”), within (“as predicted”) or lower than (“below prediction”) the 95% confidence interval of the neutral model predictions.

Normalized stochasticity ratio (NST) was used to quantify ecological stochasticity in communities within canal sections (Ning et al., 2019). This index measures the relative importance of stochasticity vs. determinism considering both the situations where deterministic factors drive the communities more similar or dissimilar than random patterns (Ning et al., 2019). An NST < 0.5 indicates the more deterministic and > 0.5 more stochastic community assembly. NST was calculated based on Jaccard similarity metrics using null model algorithm PF (fixed data richness and proportional taxa occurrence frequency). NST analysis was performed in R using the NST package (Ning et al., 2019).

#### Method S5 Population lifestyles and local growth factor analyses

The absolute abundance of bacterial and micro-eukaryotic ASVs was calculated and used to address abundance of the three neutral model partitioning groups and growth of the microbial populations in the main canal, by multiplying relative abundance of each ASV computed from amplicon sequence by the overall 16S and 18S rRNA gene copies detected in a given metagenome for all the Oct samples. For each ASV, its occurrence was computed as the proportion of samples in which the ASV was detected, and variability as the coefficient of variation of its absolute abundance across samples (Linz et al., 2017). Significant difference in occurrence and variability between bacterial and eukaryotic ASVs was tested via Welch’s *t*-test.

To characterize fate of microorganisms originated from natural reservoir after transported into the closed canal, we defined a local growth factor (LGF) to depict whether the populations grew or restrained, by comparing their mean absolute abundance in each canal section to the abundance at the canal head, as follows:

$L\mathrm{GF}_{x}=\log_{10}\frac{\mathrm{AA}_{x, N} +1}{\mathrm{AA}_{x, OctS02}+1}$	(2)

where AA_x, N_ is the average absolute abundance of ASV x in Section N of sampling sites and AA_x, OctS02_ is the absolute abundance of ASV x at OctS02, the canal head. OctS02 was excluded from Section1 for the LGF calculation. An average LGF > 0 across the three sections was considered to indicate growth in the canal.

To unravel occurrence and fate of human pathogens along the canal, the potentially pathogenic bacterial ASVs were identified referring to a list of potentially pathogenic genera (Table S2) summarized from Woolhouse et al. (2015) which presented 538 potentially human pathogenic species belonging to 141 genera, and LGFs of these identified ASVs were examined.

### Supplementary Figures

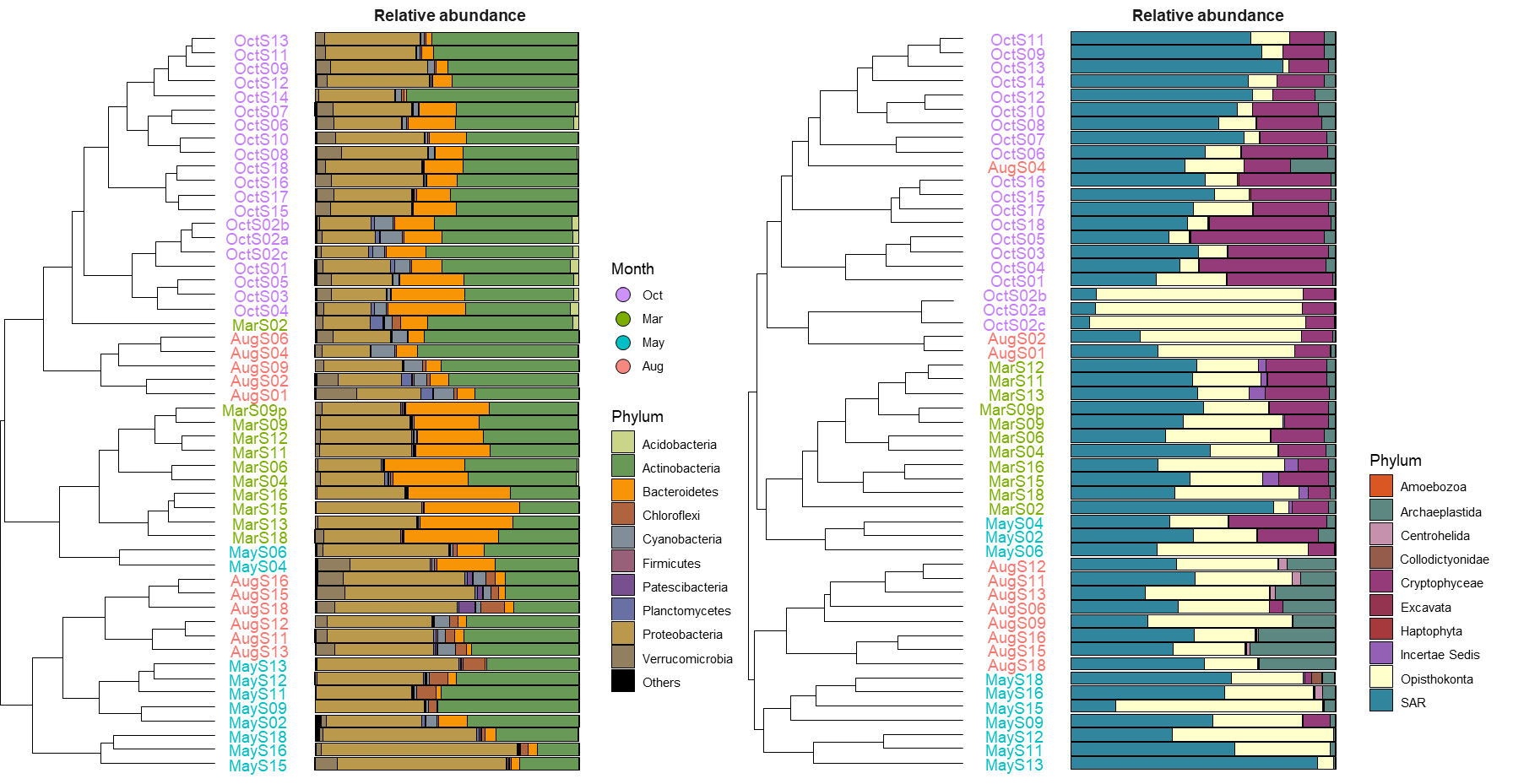

**Fig. S1** Bray-Curtis similarity-based dendrogram showing bacterial (A) and micro-eukaryotic (B) community composition for all the four sampling campaigns. Sample names are colored according to month of sampling. Relative abundance of the top 10 abundant phyla for bacteria and all the phyla for eukaryotes were shown in bar plots.

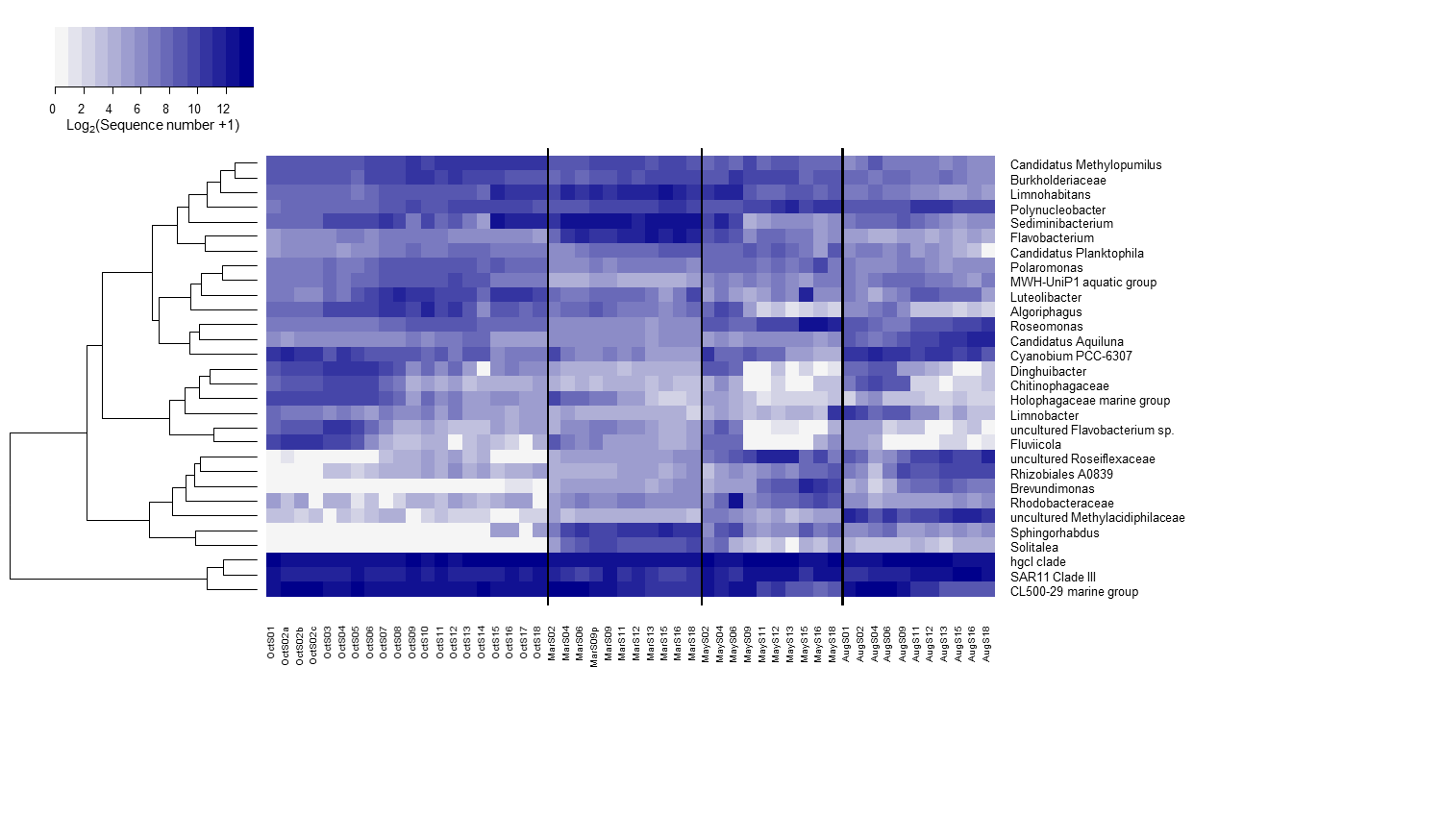

**Fig. S2** The distribution of top 30 abundant bacterial genera across the canal.

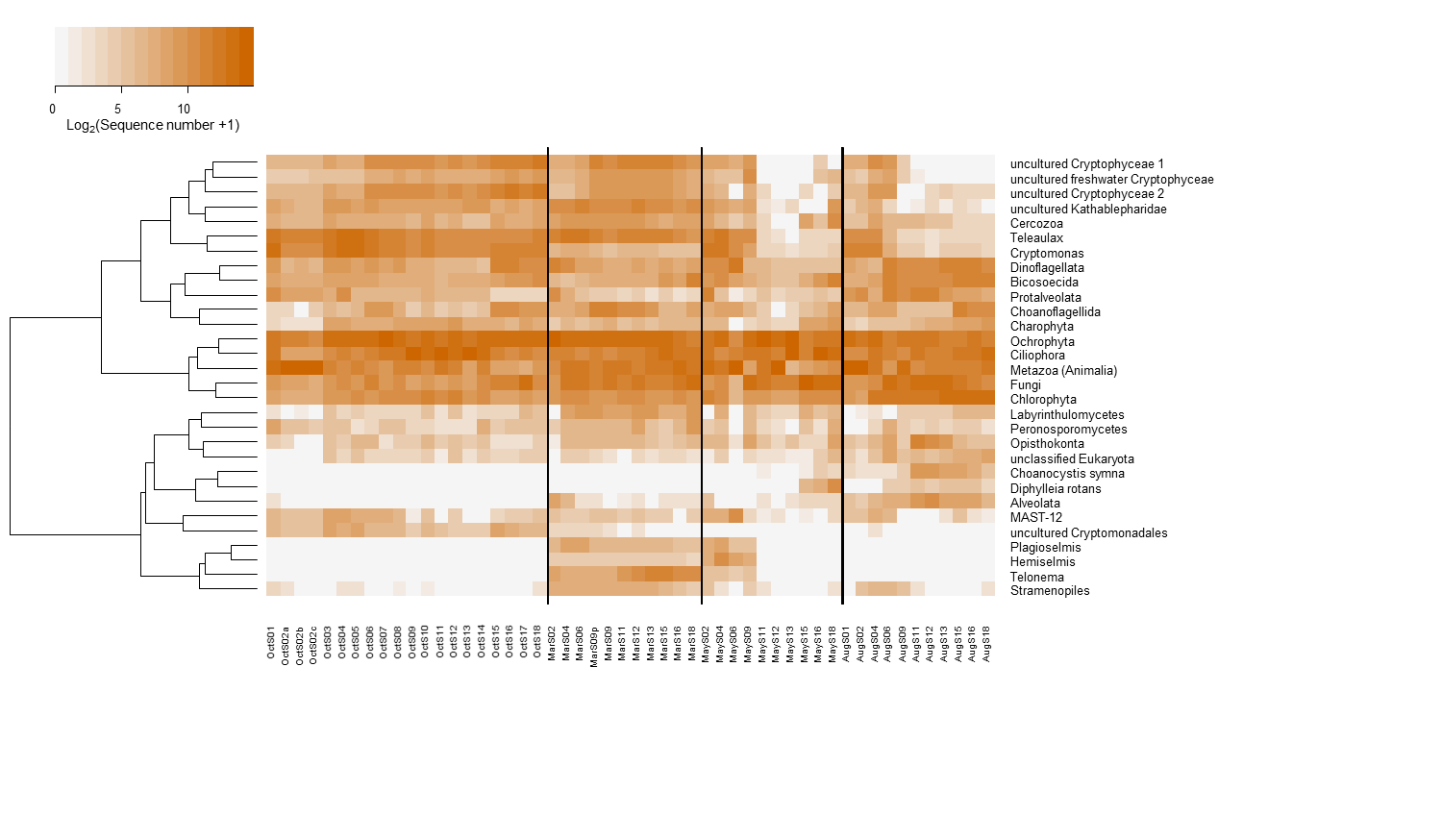

**Fig. S3** The distribution of top 30 abundant micro-eukaryotic order-level lineages across the canal.

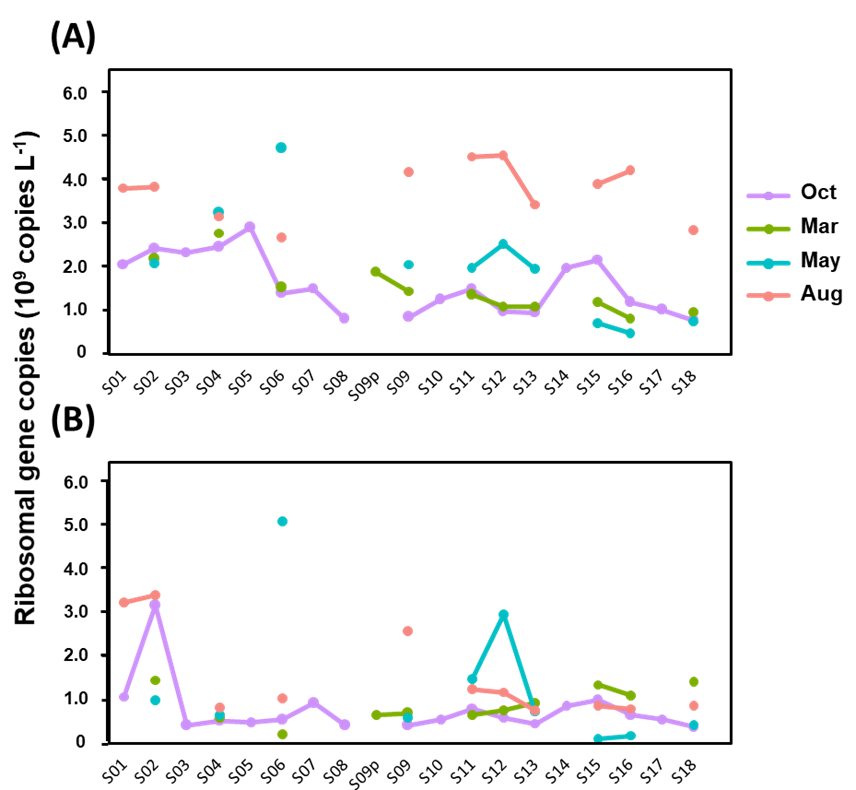

**Fig. S4** Bacterial 16S (A) and eukaryotic 18S (B) rRNA gene copies quantified for microbial communities along the canal across the four sampling campaigns.

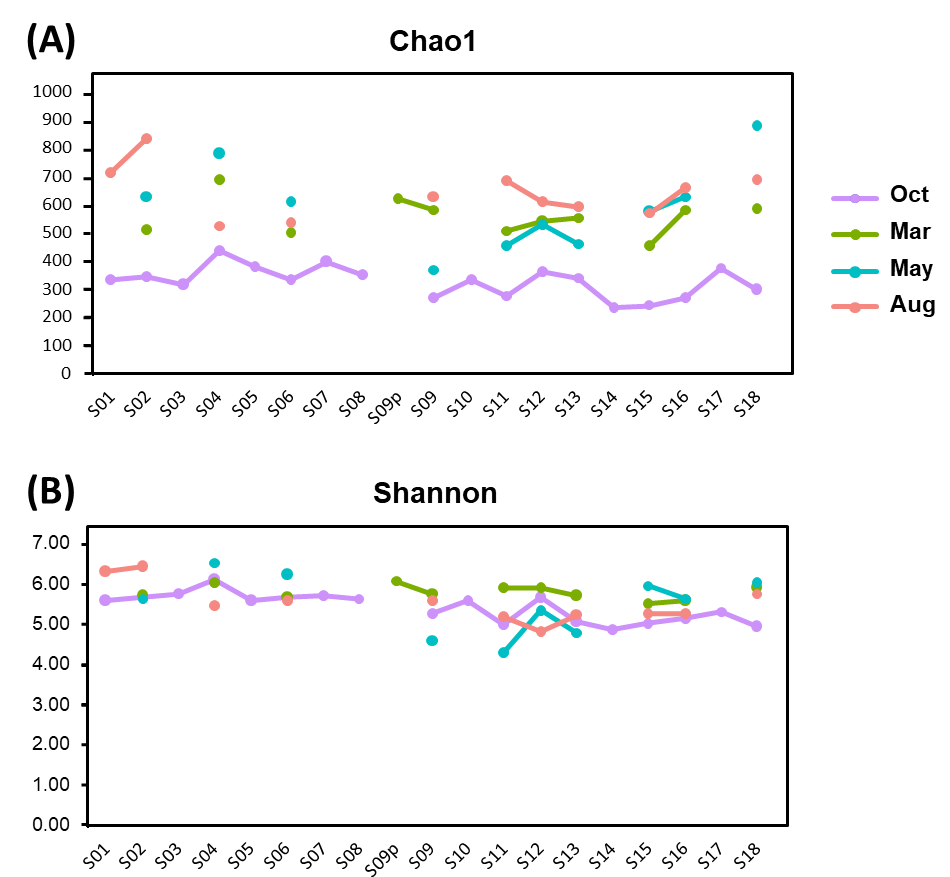

**Fig. S5** The Chao1 and Shannon indices of bacterial communities along the canal across the four samplings.

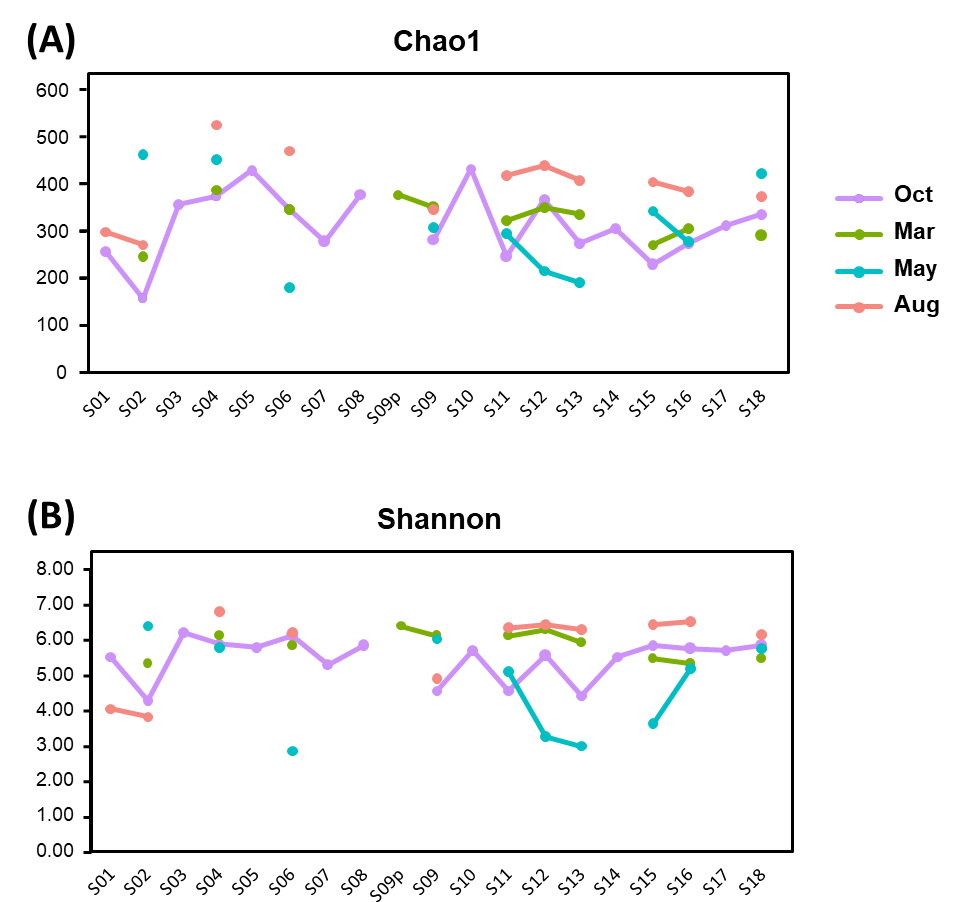

**Fig. S6** The Chao1 richness and Shannon diversity index of micro-eukaryotic communities along the canal across four samplings.

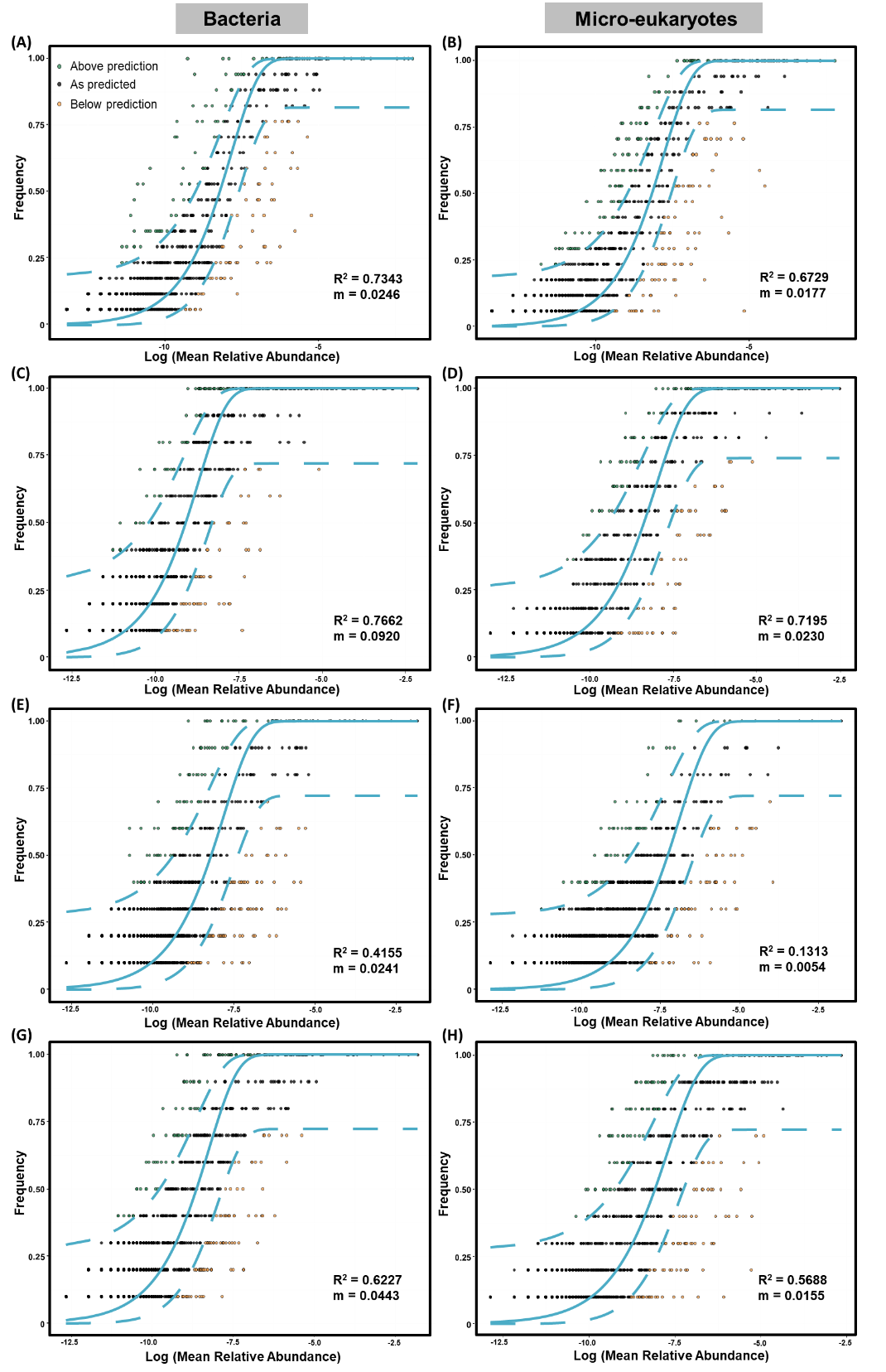

**Fig. S7** Fit of the neutral model to Oct (A, B), Mar (C, D), May (E, F) and Aug (G, H) samples across the canal for bacterial and micro-eukaryotic communities. The solid blue lines indicate the prediction of ASV occurrence frequencies predicted by the model, and the dashed blue lines represent 95% confidence intervals. ASVs that occur more frequently, less frequently than predicted or as predicted by the model are shown in different colors. Parameter “m” represents estimated migration rate.

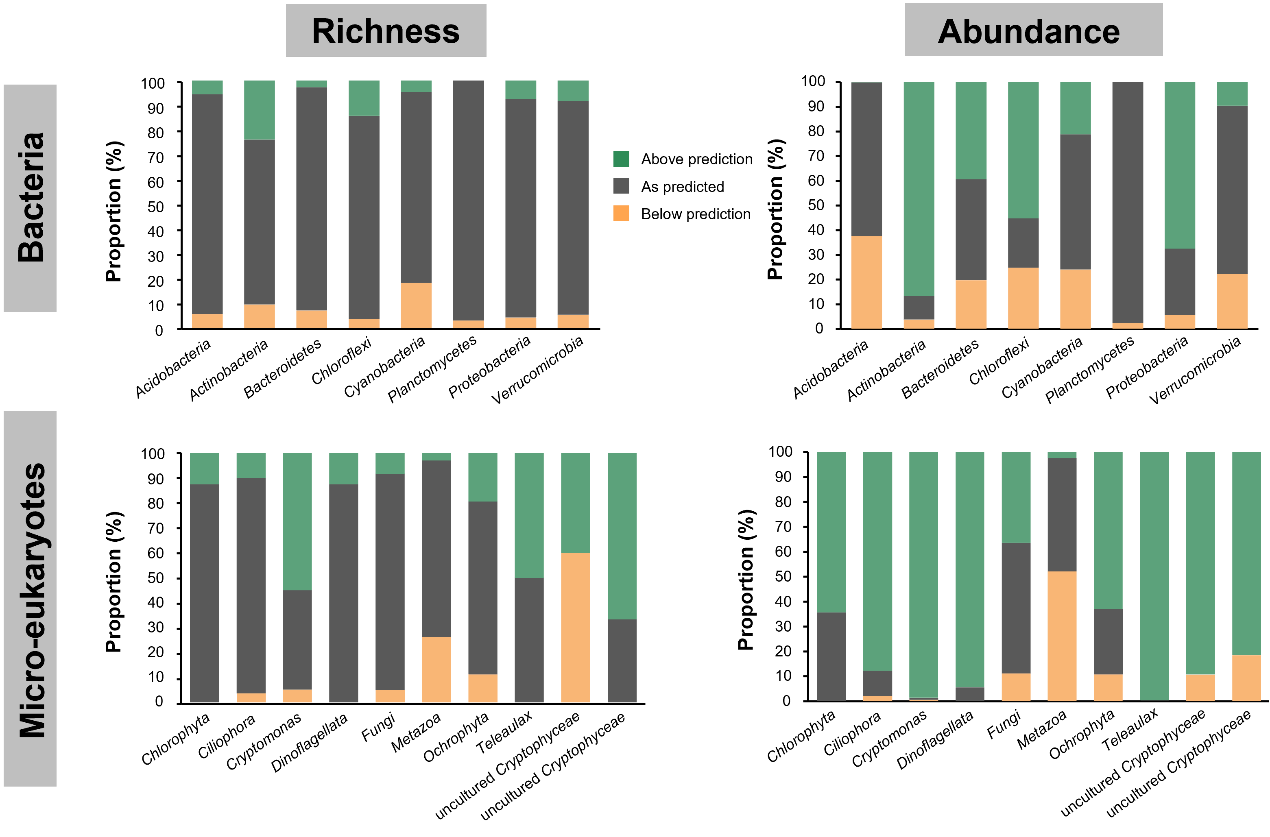

**Fig. S8** Contrasting taxa-resolved proportions of the three fitting partitions (occurrence frequency above prediction, as predicted and below prediction by the Sloan neutral model) in richness (ASV number) and abundance (sequence number) for bacterial and micro-eukaryotic communities. Only abundant bacterial phyla (> 1%) and 10 most abundant micro-eukaryotic orders were shown.

**Table S1** Ratio of bacterial 16S and eukaryotic 18S rRNA gene copies along the canal based on metagenomic sequencing of the canal water samples.

| **Sample** | **16S/18S** | **No. of 16S reads** | **No. of 18S reads** |
| --- | --- | --- | --- |
| OcS01 | 1.96 | 5751 | 2928 |
| OcS02 | 0.76 | 5806 | 7639 |
| OcS03 | 5.77 | 7864 | 1363 |
| OcS04 | 4.78 | 6207 | 1299 |
| OcS05 | 6.24 | 7729 | 1238 |
| OcS06 | 2.64 | 5367 | 2036 |
| OcS07 | 1.64 | 5416 | 3312 |
| OcS08 | 2.11 | 4800 | 2273 |
| OcS09 | 2.14 | 6086 | 2843 |
| OcS10 | 2.26 | 5482 | 2423 |
| OcS11 | 1.91 | 5501 | 2876 |
| OcS12 | 1.77 | 5708 | 3228 |
| OcS13 | 2.21 | 4947 | 2240 |
| OcS14 | 2.36 | 4808 | 2037 |
| OcS15 | 2.15 | 4066 | 1888 |
| OcS16 | 1.87 | 4813 | 2577 |
| OcS17 | 1.93 | 4825 | 2502 |
| OcS18 | 2.18 | 3807 | 1747 |
| MarS02 | 1.54 | 7050 | 4583 |
| MarS04 | 4.71 | 7839 | 1665 |
| MarS06 | 7.07 | 7631 | 1079 |
| MarS09p | 2.75 | 7203 | 2617 |
| MarS09 | 2.22 | 6683 | 3014 |
| MarS11 | 2.06 | 6572 | 3195 |
| MarS12 | 1.46 | 6173 | 4226 |
| MarS13 | 1.18 | 5822 | 4930 |
| MarS15 | 0.91 | 5382 | 5919 |
| MarS16 | 0.73 | 5026 | 6869 |
| MarS18 | 0.68 | 6923 | 10164 |
| MayS02 | 2.14 | 7384 | 3456 |
| MayS04 | 5.00 | 7397 | 1480 |
| MayS06 | 0.93 | 6359 | 6806 |
| MayS09 | 3.53 | 6820 | 1930 |

**Table S1** continued

| **Sample** | **16S/18S** | **No. of 16S reads** | **No. of 18S reads** |
| --- | --- | --- | --- |
| MayS11 | 1.33 | 6459 | 4852 |
| MayS12 | 0.86 | 6259 | 7301 |
| MayS13 | 2.74 | 4712 | 1720 |
| MayS15 | 6.29 | 3735 | 594 |
| MayS16 | 3.15 | 4685 | 1487 |
| MayS18 | 1.81 | 5411 | 2987 |
| AugS01 | 1.18 | 5242 | 4450 |
| AugS02 | 1.13 | 5085 | 4482 |
| AugS04 | 3.90 | 5041 | 1292 |
| AugS06 | 2.58 | 4994 | 1937 |
| AugS09 | 1.63 | 5467 | 3361 |
| AugS11 | 3.64 | 5769 | 1587 |
| AugS12 | 3.89 | 6100 | 1570 |
| AugS13 | 4.58 | 5128 | 1119 |
| AugS15 | 4.65 | 6178 | 1330 |
| AugS16 | 5.40 | 4852 | 899 |
| AugS18 | 3.40 | 6043 | 1779 |
| MayS11 | 1.33 | 6459 | 4852 |

**Table S2** List of potentially pathogenic genera summarized according to Woolhouse et al. (2015). Genera highlighted in red were detected in this study.

| *Abiotrophia* | *Borrelia* | *Delftia* | *Gordonia* | *Mycobacterium* | *Providencia* | *Stenotrophomonas* |
| --- | --- | --- | --- | --- | --- | --- |
| *Achromobacter* | *Brevibacillus* | *Dermatophilus* | *Granulicatella* | *Mycoplasma* | *Pseudomonas* | *Streptobacillus* |
| *Acidaminococcus* | *Brevundimonas* | *Dichelobacter* | *Haemophilus* | *Myroides* | *Pseudonocardia* | *Streptococcus* |
| *Acinetobacter* | *Brucella* | *Edwardsiella* | *Hafnia* | *Neisseria* | *Pseudoramibacter* | *Sutterella* |
| *Actinobacillus* | *Burkholderia* | *Eggerthella* | *Helicobacter* | *Neorickettsia* | *Psychrobacter* | *Suttonella* |
| *Actinomyces* | *Campylobacter* | *Ehrlichia* | *Kingella* | *Nocardia* | *Rahnella* | *Tatlockia* |
| *Aerococcus* | *Capnocytophaga* | *Eikenella* | *Klebsiella* | *Ochrobactrum* | *Ralstonia* | *Tatumella* |
| *Aeromonas* | *Cardiobacterium* | *Enterobacter* | *Kluyvera* | *Oligella* | *Rhodococcus* | *Treponema* |
| *Alcaligenes* | *Cedecea* | *Enterococcus* | *Lactobacillus* | *Orientia* | *Rickettsia* | *Tropheryma* |
| *Amycolatopsis* | *Cellulomonas* | *Erysipelothrix* | *Legionella* | *Paenibacillus* | *Rothia* | *Tsukamurella* |
| *Anaerococcus* | *Centipeda* | *Escherichia* | *Leifsonia* | *Pantoea* | *Ruminococcus* | *Ureaplasma* |
| *Arcanobacterium* | *Chlamydia* | *Eubacterium* | *Leptospira* | *Pasteurella* | *Saccharomonospora* | *Veillonella* |
| *Arcobacter* | *Chlamydophila* | *Ewingella* | *Leptotrichia* | *Peptinophilus* | *Saccharopolyspora* | *Vibrio* |
| *Anaplasma* | *Chromobacterium* | *Fibrobacter* | *Listeria* | *Peptococcus* | *Salmonella* | *Wolinella* |
| *Bacillus* | *Chryseobacterium* | *Filifactor* | *Mannheimia* | *Peptostreptococcus* | *Sebaldella* | *Yersinia* |
| *Bacteroides* | *Citrobacter* | *Finegoldia* | *Megamonas* | *Photobacterium* | *Selenomonas* |  |
| *Bartonella* | *Clostridium* | *Fluoribacter* | *Megasphaera* | *Plesiomonas* | *Serratia* |  |
| *Bergeyella* | *Collinsella* | *Francisella* | *Micromonas* | *Porphyromonas* | *Shigella* |  |
| *Bifidobacterium* | *Comamonas* | *Fusobacterium* | *Mogibacterium* | *Prevotella* | *Sphingomonas* |  |
| *Bilophila* | *Corynebacterium* | *Gardnerella* | *Moraxella* | *Propionibacterium* | *Spirillum* |  |
| *Bordetella* | *Coxiella* | *Gemella* | *Morganella* | *Proteus* | *Staphylococcus* |  |

**Table S3** Physiochemical properties of water samples collected from 16 sites over four seasons.

| **Sample** | **T (°C)** | **pH** | **DO (mg L^-1^)** | **TN (mg L^-1^)** | **F^-^ (mg L^-1^)** |
| --- | --- | --- | --- | --- | --- |
| OctS02 | 20.5 | 8.3 | 8.9 | 1.095 | 0.167 |
| OctS03 | 20.5 | 8.3 | 9.1 | 1.140 | 0.165 |
| OctS04 | 19.8 | 8.5 | 10.9 | 1.240 | 0.203 |
| OctS05 | 20.0 | 8.4 | 11.1 | 1.175 | 0.173 |
| OctS06 | 19.7 | 8.5 | 9.9 | 1.125 | 0.188 |
| OctS07 | 19.6 | 8.5 | 11.4 | 1.165 | 0.163 |
| OctS08 | 18.8 | 8.5 | 10.4 | 1.035 | 0.170 |
| OctS09 | 18.7 | 8.6 | 11.1 | 1.010 | 0.154 |
| OctS10 | 18.9 | 8.3 | 8.9 | 1.245 | 0.160 |
| OctS12 | 17.5 | 8.3 | 9.0 | 1.285 | 0.145 |
| OctS13 | 18.5 | 8.4 | 9.4 | 1.215 | 0.150 |
| OctS14 | 18.3 | 8.5 | 9.2 | 1.210 | 0.155 |
| OctS15 | 12.1 | 8.2 | 10.3 | 1.190 | 0.175 |
| OctS16 | 10.1 | 8.2 | 11.0 | 1.175 | 0.185 |
| OctS17 | 11.4 | 8.3 | 10.3 | 1.090 | 0.175 |
| OctS18 | 12.7 | 8.3 | 11.1 | 1.355 | 0.185 |
| MarS02 | 8.4 | 8.1 | 9.8 | 1.170 | 0.187 |
| MarS04 | 9.1 | 8.0 | 9.6 | 1.080 | 0.188 |
| MarS06 | 8.9 | 8.3 | 9.2 | 1.030 | 0.194 |
| MarS09 | 9.0 | 8.2 | 8.8 | 1.110 | 0.203 |
| MarS12 | 9.5 | 8.2 | 9.5 | 1.270 | 0.170 |
| MarS13 | 9.5 | 8.2 | 10.8 | 1.260 | 0.200 |
| MarS15 | 7.1 | 8.0 | 10.9 | 1.210 | 0.210 |
| MarS16 | 8.0 | 8.0 | 10.7 | 1.180 | 0.220 |
| MarS18 | 5.3 | 8.0 | 12.2 | 1.100 | 0.220 |
| MayS02 | 16.9 | 8.0 | 8.4 | 1.030 | 0.221 |
| MayS04 | 18.9 | 8.1 | 8.3 | 1.070 | 0.213 |
| MayS06 | 18.0 | 8.2 | 9.0 | 1.100 | 0.205 |
| MayS09 | 19.9 | 8.2 | 9.0 | 1.020 | 0.205 |
| MayS12 | 19.2 | 8.3 | 8.7 | 1.030 | 0.210 |
| MayS13 | 21.0 | 8.4 | 9.0 | 1.150 | 0.220 |
| MayS15 | 20.0 | 8.1 | 9.8 | 1.230 | 0.160 |
| MayS16 | 19.5 | 8.1 | 9.7 | 1.020 | 0.160 |
| MayS18 | 19.2 | 8.0 | 10.9 | 1.060 | 0.230 |
| AugS02 | 28.2 | 7.8 | 7.5 | 1.000 | 0.146 |
| AugS04 | 26.8 | 8.0 | 8.2 | 0.940 | 0.159 |
| AugS06 | 29.0 | 8.1 | 8.4 | 0.930 | 0.153 |
| AugS09 | 28.9 | 8.4 | 9.2 | 0.950 | 0.168 |
| AugS12 | 30.0 | 8.4 | 7.9 | 0.910 | 0.180 |
| AugS13 | 29.6 | 8.3 | 7.9 | 0.830 | 0.170 |
| AugS15 | 30.2 | 8.3 | 9.0 | 0.850 | 0.160 |
| AugS16 | 29.6 | 8.3 | 8.9 | 0.860 | 0.160 |
| AugS18 | 30.8 | 8.2 | 8.4 | 1.240 | 0.200 |

**Table S4** PCNM (Principal Coordinates of Neighborhood Matrices) of the 16 sites where water properties were measured.

| **Sample** | **PCNM1** | **PCNM2** | **PCNM3** | **PCNM4** | **PCNM5** | **PCNM6** | **PCNM7** | **PCNM8** | **PCNM9** | **PCNM10** |
| --- | --- | --- | --- | --- | --- | --- | --- | --- | --- | --- |
| OcS02 | -0.1223 | 0.1599 | -0.2297 | 0.2663 | -0.3125 | 0.3071 | 0.3327 | 0.3121 | -0.3308 | 0.2957 |
| OcS03 | -0.2272 | 0.2551 | -0.3381 | 0.2928 | -0.2771 | 0.1326 | 0.0655 | -0.0962 | 0.1785 | -0.2957 |
| OcS04 | -0.3068 | 0.2652 | -0.2734 | 0.0601 | 0.0681 | -0.2793 | -0.3252 | -0.3422 | 0.2358 | -0.0744 |
| OcS05 | -0.3438 | 0.1847 | -0.0703 | -0.2427 | 0.3264 | -0.3252 | -0.1232 | 0.1137 | -0.3017 | 0.3139 |
| OcS06 | -0.3302 | 0.0291 | 0.1744 | -0.3925 | 0.2200 | 0.0575 | 0.3092 | 0.2876 | -0.0769 | -0.2373 |
| OcS07 | -0.2728 | -0.1552 | 0.3261 | -0.2716 | -0.1311 | 0.3151 | 0.1838 | -0.2052 | 0.3404 | -0.1410 |
| OcS08 | -0.1799 | -0.3228 | 0.3086 | 0.0274 | -0.3405 | 0.0806 | -0.2753 | -0.2945 | -0.1130 | 0.3283 |
| OcS09 | -0.0642 | -0.4203 | 0.1293 | 0.2650 | -0.1699 | -0.3026 | -0.2294 | 0.2122 | -0.2728 | -0.1824 |
| OcS10 | 0.0632 | -0.4182 | -0.1227 | 0.2617 | 0.1933 | -0.2932 | 0.2355 | 0.2191 | 0.2598 | -0.2083 |
| OcS12 | 0.1819 | -0.3197 | -0.3090 | 0.0204 | 0.3392 | 0.0960 | 0.2759 | -0.2912 | 0.1293 | 0.3277 |
| OcS13 | 0.2779 | -0.1545 | -0.3332 | -0.2730 | 0.1056 | 0.3098 | -0.1835 | -0.2098 | -0.3365 | -0.1148 |
| OcS14 | 0.3331 | 0.0287 | -0.1806 | -0.3817 | -0.2409 | 0.0382 | -0.3035 | 0.2827 | 0.0523 | -0.2577 |
| OcS15 | 0.3429 | 0.1867 | 0.0727 | -0.2250 | -0.3240 | -0.3374 | 0.1308 | 0.1230 | 0.3134 | 0.3090 |
| OcS16 | 0.3069 | 0.2702 | 0.2826 | 0.0692 | -0.0481 | -0.2680 | 0.3241 | -0.3390 | -0.2232 | -0.0516 |
| OcS17 | 0.2252 | 0.2560 | 0.3398 | 0.2776 | 0.2826 | 0.1465 | -0.0679 | -0.1032 | -0.1958 | -0.3018 |
| OcS18 | 0.1161 | 0.1552 | 0.2232 | 0.2458 | 0.3087 | 0.3224 | -0.3496 | 0.3307 | 0.3412 | 0.2903 |

**Table S5** Seasonal variation of environmental variables. Significance of seasonal effects were tested with the Kruskal-Wallis one-way ANOVA.

|  | **T (°C)** | | **pH** | | **DO (mg L^-1^)** | | **TN (mg L^-1^)** | | **F^-^ (mg L^-1^)** | |
| --- | --- | --- | --- | --- | --- | --- | --- | --- | --- | --- |
|  | ***H*** | ***P*** | ***H*** | ***P*** | ***H*** | ***P*** | ***H*** | ***P*** | ***H*** | ***P*** |
| Seasonal effect | 33.39 | **< 10^-6^** | 16.67 | **< 10^-3^** | 18.73 | **< 10^-3^** | 18.22 | **< 10^-3^** | 18.42 | **< 10^-3^** |

**Table S6** Bacterial ASVs with significant and strong correlation with fluoride. Correlation was tested by computing Spearman’s rank correlation, corrected with the Benjamini-Hochberg method. Only ASVs with Spearman’s r_s_ > 0.6 or < -0.6, and *P* < 0.05 are shown.

|  | **Spearman’s r_s_** | ***P*** | **Taxonomy** | | | | | | |
| --- | --- | --- | --- | --- | --- | --- | --- | --- | --- |
|  |  |  | **Kingdom** | **Phylum** | **Class** | | **Order** | **Family** | **Genus** |
| ASV75 | 0.71 | < 0.001 | Bacteria | Proteobacteria | | Gammaproteobacteria | Betaproteobacteriales | Burkholderiaceae | NA |
| ASV316 | 0.68 | < 0.01 | Bacteria | Actinobacteria | | Actinobacteria | Frankiales | Sporichthyaceae | Candidatus Planktophila |
| ASV128 | 0.64 | < 0.01 | Bacteria | Proteobacteria | | Gammaproteobacteria | Betaproteobacteriales | Burkholderiaceae | Rhodoferax |
| ASV113 | 0.63 | < 0.01 | Bacteria | Actinobacteria | | Actinobacteria | Frankiales | Sporichthyaceae | hgcI clade |
| ASV156 | -0.71 | < 0.001 | Bacteria | Proteobacteria | | Alphaproteobacteria | Rickettsiales | Rickettsiaceae | Candidatus Megaira |
| ASV48 | -0.68 | < 0.01 | Bacteria | Actinobacteria | | Acidimicrobiia | Microtrichales | Ilumatobacteraceae | CL500-29 marine group |
| ASV108 | -0.65 | < 0.01 | Bacteria | Proteobacteria | | Alphaproteobacteria | Rhizobiales | Beijerinckiaceae | FukuN57 |
| ASV147 | -0.64 | < 0.01 | Bacteria | Cyanobacteria | | Oxyphotobacteria | Nostocales | Microcystaceae | Microcystis PCC-7914 |
| ASV193 | -0.60 | < 0.05 | Bacteria | Proteobacteria | | Gammaproteobacteria | Gammaproteobacteria Incertae Sedis | Unknown Family | Acidibacter |
| ASV430 | -0.60 | < 0.05 | Bacteria | Proteobacteria | | Alphaproteobacteria | Rickettsiales | AB1 | uncultured bacterium |
| ASV76 | -0.60 | < 0.05 | Bacteria | Proteobacteria | | Gammaproteobacteria | Betaproteobacteriales | Burkholderiaceae | MWH-UniP1 aquatic group |

**Table S7** Micro-eukaryotic ASVs with significant and strong correlation with fluoride. Correlation was tested by computing Spearman’s rank correlation, corrected with the Benjamini-Hochberg method. Only ASVs with Spearman’s r_s_ > 0.6 or < -0.6, and *P* < 0.05 are shown.

|  | **Spearman’s r_s_** | | ***P*** | **Taxonomy** | | | | | |
| --- | --- | --- | --- | --- | --- | --- | --- | --- | --- |
|  |  |  |  | **Kingdom** | **Phylum** | **Class** | **Order** | **Family** | **Genus** |
| ASV169 | | 0.66 | < 0.01 | Eukaryota | SAR | Stramenopiles | Ochrophyta | Ochromonadales | uncultured |
| ASV91 | | 0.63 | < 0.01 | Eukaryota | Archaeplastida | Chloroplastida | Chlorophyta | Sphaeropleales | Mychonastes |
| ASV30 | | 0.62 | < 0.01 | Eukaryota | SAR | Stramenopiles | Ochrophyta | Chrysophyceae | P34.48 |
| ASV157 | | 0.61 | < 0.01 | Eukaryota | SAR | Stramenopiles | Bicosoecida | LG08-10 | uncultured freshwater eukaryote |
| ASV273 | | 0.60 | < 0.01 | Eukaryota | SAR | Stramenopiles | Ochrophyta | Chrysophyceae | uncultured |
| ASV298 | | 0.60 | < 0.01 | Eukaryota | SAR | Alveolata | Dinoflagellata | Suessiaceae | Symbiodinium |
| ASV26 | | -0.68 | < 0.001 | Eukaryota | SAR | Stramenopiles | Ochrophyta | Mediophyceae | NA |
| ASV431 | | -0.68 | < 0.001 | Eukaryota | Archaeplastida | Chloroplastida | Chlorophyta | Chlamydomonadales | NA |
| ASV19 | | -0.63 | < 0.01 | Eukaryota | SAR | Stramenopiles | Ochrophyta | Mediophyceae | Discostella |
| ASV56 | | -0.64 | < 0.01 | Eukaryota | Archaeplastida | Chloroplastida | Chlorophyta | Chlorophyceae | NA |
| ASV6 | | -0.64 | < 0.01 | Eukaryota | Opisthokonta | Holozoa | Metazoa (Animalia) | Copepoda | Calanoida |
| ASV170 | | -0.61 | < 0.01 | Eukaryota | SAR | Alveolata | Protalveolata | Perkinsidae | A31 |
| ASV180 | | -0.60 | < 0.01 | Eukaryota | Archaeplastida | Chloroplastida | Chlorophyta | Sphaeropleales | Ambiguous_taxa |
| ASV39 | | -0.61 | < 0.01 | Eukaryota | SAR | Alveolata | Ciliophora | Phyllopharyngea | Suctoria |
| ASV69 | | -0.60 | < 0.01 | Eukaryota | Archaeplastida | Chloroplastida | Chlorophyta | Sphaeropleales | Radiococcus |

**Table S8** Correlation between ribosomal RNA gene copies, diversity metrics and environmental variables.

|  | T (°C) | | pH | | DO (mg L^-1^) | | TN (mg L^-1^) | | F^-^ (mg L^-1^) | |
| --- | --- | --- | --- | --- | --- | --- | --- | --- | --- | --- |
|  | Spearman’s r_s_ | *P* | Spearman’s r_s_ | *P* | Spearman’s r_s_ | *P* | Spearman’s r_s_ | *P* | Spearman’s r_s_ | *P* |
| 16S copies (copies L^-1^) | 0.55 | **< 0.01** | 0.05 | 0.74 | -0.61 | **< 0.01** | -0.48 | **< 0.01** | -0.05 | 0.76 |
| 16S Chao1 richness | 0.20 | 0.20 | -0.52 | **< 0.01** | -0.38 | **0.01** | -0.42 | **< 0.01** | 0.30 | 0.05 |
| 16S Shannon’s diversity index | -0.15 | 0.33 | -0.38 | **0.01** | 0.03 | 0.87 | 0.16 | 0.31 | 0.17 | 0.27 |
| 18S copies (copies L^-1^) | 0.04 | 0.81 | -0.26 | 0.10 | -0.32 | **0.03** | -0.19 | 0.22 | 0.16 | 0.30 |
| 18S Chao1 richness | 0.33 | **0.03** | -0.04 | 0.79 | -0.31 | **0.04** | -0.25 | 0.11 | -0.08 | 0.61 |
| 18S Shannon’s diversity index | 0.21 | 0.17 | -0.07 | 0.66 | -0.27 | 0.08 | -0.23 | 0.14 | 0.01 | 0.95 |
